# ExEnDiff: An Experiment-guided Diffusion model for protein conformational Ensemble generation

**DOI:** 10.1101/2024.10.04.616517

**Authors:** Yikai Liu, Zongxin Yu, Richard J. Lindsay, Guang Lin, Ming Chen, Abhilash Sahoo, Sonya M. Hanson

**Affiliations:** Department of Mechanical Engineering, Purdue University, West Lafayette, IN, 47907; Center for Computational Mathematics, Flatiron Institute, New York, NY, USA; Center for Computational Biology, Flatiron Institute, New York, NY, USA; Department of Chemistry, Purdue University, West Lafayette, IN, 47907; Department of Engineering Sciences and Applied Math, Northwestern University, Evanston, IL, 60201

## Abstract

Understanding protein conformation is key to understanding their function. Importantly, most proteins adopt multiple conformations with non-trivial ensemble distributions that change depending on their environment to perform functions like catalysis, signaling, and transport. Recently, machine learning techniques, especially deep generative models, have been employed to develop protein conformation generators. These models, known as unified protein ensemble samplers, are trained on the PDB dataset and can generate diverse protein conformation ensembles given a protein sequence. However, their reliance solely on structural data from the PDB, which primarily captures folded protein states, restricts the diversity of the generated ensembles and can result in physically unrealistic conformations. In this paper, we overcome these challenges by introducing ExEnDiff, an experiment-guided diffusion model for protein conformation generation. ExEnDiff integrates experimental measurements as a physical prior, enabling the generation of protein conformations with desired properties. Our experiments on a variety of fast-folding and intrinsically disordered proteins demonstrate that ExEnDiff significantly advances the capabilities of current unified protein ensemble samplers. With little computational cost, ExEnDiff can capture important proteins’ configuration properties and the underlying Boltzmann distribution, paving the way for a next-generation molecular dynamics engine. We further demonstrate the effectiveness of ExEnDiff to capture conformational changes in the presence of mutations and as an efficient tool for determining a reasonable CV space for protein ensembles. With these results, ExEnDiff is well-poised to push the study of protein ensembles into a data-rich regime currently available to few problems in biology.

## Introduction

The dynamic nature of biological macromolecules is fundamental to their capacity to perform essential functions within the cell. For proteins in particular, these conformational changes are crucial because they regulate protein activity and dictate interactions with other biomolecules, allowing precise control in complex physiological processes such as signaling, transport, and gene expression. Understanding this conformational flexibility is therefore essential for comprehending how proteins function in both normal physiological states and pathological conditions, offering insights that are critical for advancing therapeutic development. However, our ability to capture this conformational variability is hindered by the limitations of current experimental and computational methods. Experimental techniques like NMR and cryo-EM are resource-intensive and often miss transient states, while computational approaches like molecular dynamics require significant computational power to explore relevant timescales.

Recent advancements in artificial intelligence (AI) have transformed structural biology, particularly within the problem space of determining a folded structure from amino acid sequence alone. AI-based approaches to this problem, such as AlphaFold, ^1,2^ RoseTTAFold, ^3^ trRosetta, ^4^ and ESMFold,^5^ have significantly enhanced our ability to predict and analyze protein conformations with unprecedented accuracy. More recent efforts have extended beyond single structure prediction to generate protein conformational ensembles. Initial methods, such as MSA sub-sampling^6^ and clustering,^7^ have attempted to diversify AlphaFold’s predicted conformations by adjusting model inputs. More advanced strategies have focused on deep generative models, particularly diffusion models, ^8,9^ to address the challenge of generating protein conformation ensembles.^10–13^ These models operate on the premise that metastable states of proteins are encoded within the native structures of other proteins. By training on large datasets of native protein structures from the Protein Data Bank (PDB), these models use a stochastic perturbation process to explore conformation space with greater diversity and accuracy. This emerging class of deep generative models, referred to as unified protein ensemble samplers, shows great promise as an avenue for generating protein ensembles.

However, PDB predominantly contains experimentally resolved, folded protein structures. This limits the ability of the database to capture the full spectrum of protein conformations, particularly in less-structured regions, Thus, unified protein ensemble samplers trained solely on PDB data struggle to accurately explore the entire conformational landscape of proteins. At the time of AlphaFold2, the PDB had ∼140,000 well-curated structures (and evolutionary couplings were trained on a database of of more than two billion clustered protein sequences) providing a wealth of training data to learn static structures. For protein ensembles, this wealth of data is not available. Furthermore much of the experimental data available for ensemble properties of proteins are ensemble measurements that are difficult to connect to a molecular representation of the ensemble. Computational methods like molecular dynamics (MD) simulation can provide access to protein ensembles,^14–17^ but are computationally intensive, especially for large proteins. ^18^ Efforts to integrate physical knowledge, such as using MD energy as a prior, ^19^ have been made to align sampled conformations with the underlying Boltzmann distribution. However, these approaches often require costly retraining and are constrained to implicit solvent force fields, limiting their accuracy and broader applicability.

In this study, we introduce an experiment-guided diffusion model for protein conformation generation (ExEnDiff), a novel deep generative unified protein ensemble sampler that leverages experimental measurements to guide the protein conformation sampling process. ExEnDiff seamlessly integrates experimental data without incurring additional computational overhead and is compatible with any state-of-the-art deep generative unified protein ensemble sampler. The core innovation of ExEnDiff lies in its ability to direct the sampling process in alignment with experimental measurements, ensuring that the generated ensembles closely adhere to desired physical properties and accurately reflect the underlying Boltzmann distribution.

We demonstrate the efficacy of ExEnDiff through extensive protein conformation generation experiments on a range of fast-folding and intrinsically disordered proteins, achieving high-quality sampling without requiring model retraining. Numerical evaluations show that ExEnDiff consistently outperforms a state-of-the-art (SOTA) unified protein ensemble sampler, especially for proteins with flexible and disordered regions. We propose that ExEnDiff serves as a versatile computational tool for rapidly generating protein ensembles that closely follow the Boltzmann distribution, with broad applications in tasks such as collective variable selection, force field parameterization, and high-throughput screening.

## Results

### Overview of the ExEnDiff Model

The high-level methodology framework of ExEnDiff is illustrated in Fig. 1. Similar to other unified protein ensemble samplers, ExEnDiff leverages a diffusion model, as depicted in Fig. 1(a). The diffusion model perturbs the data distribution toward a simple gaussian distribution using a stochastic differential equation (SDE), and then learns the score function in the reverse SDE to recover the original data distribution. Unified protein ensemble samplers utilize the stochasticity of the diffusion model to generate diverse protein conformation ensembles through multiple steps of the reverse diffusion process. However, without guidance, these samplers may produce ensembles that significantly deviate from the true Boltzmann distribution. Inspired by restrained ensemble models^20–22^ and experiment-guided MD simulations, ExEnDiff addresses this limitation by incorporating experimentally measured observables into the sampling process as a correction term, as shown in Fig. 1(b). During the reverse diffusion process, these experimental measurements guide the sampler to align closer to the true experimental values, as depicted in Fig. 1(c). As a result, ExEnDiff enables the unified protein ensemble sampler to capture both critical global and local features of protein ensembles, generating conformations that more accurately follow the Boltzmann distribution compared to guide-free samplers. Formulations of the diffusion model and ExEnDiff are detailed in Method section.

**Figure 1.**
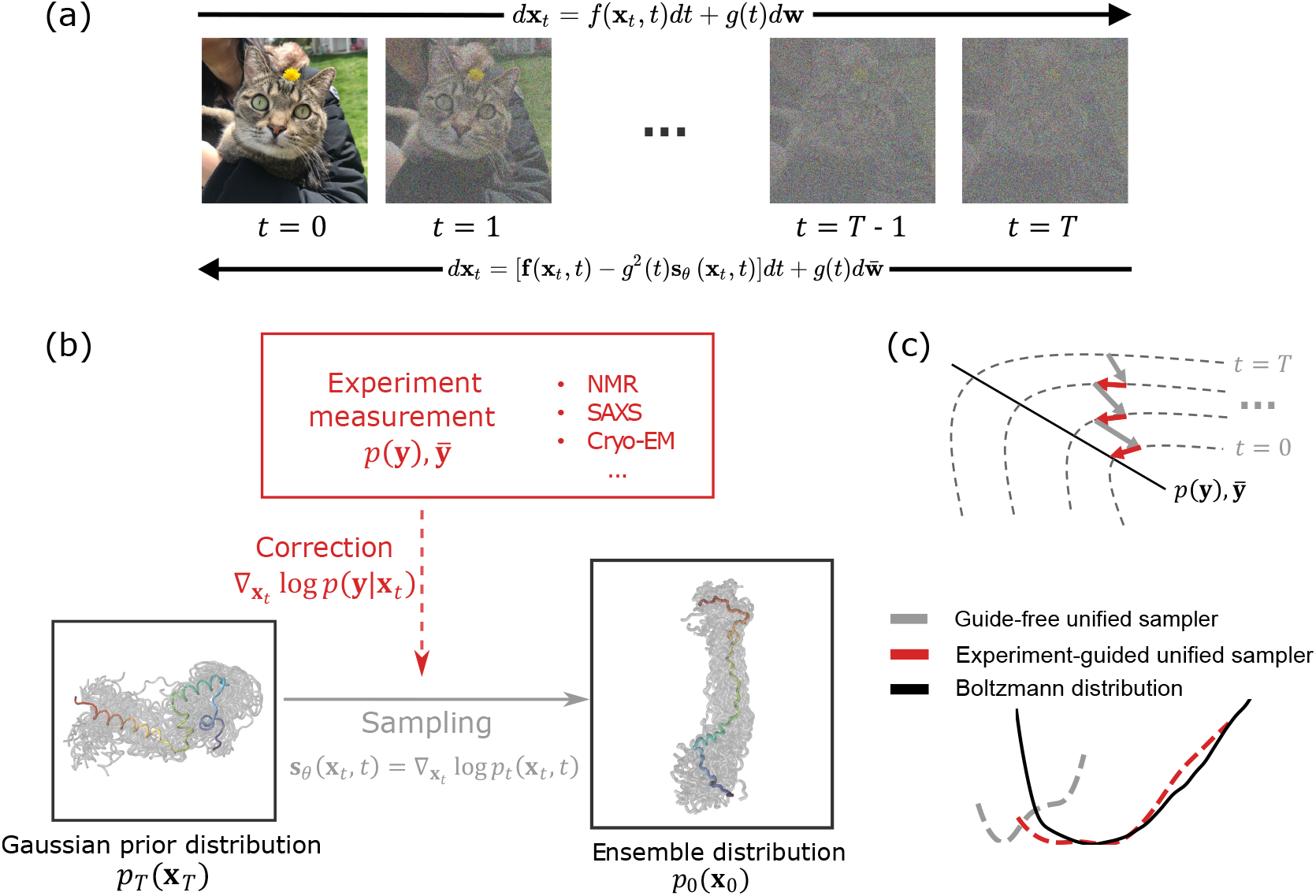
The high level framework of ExEnDiff. (a) A visual demonstration of score-based diffusion model: perturb the data distribution to a simple prior distribution through a forward stochastic differential equation and learn the reverse diffusion process to recover the correct data distribution. (b) Unified protein ensemble samplers generate initial samples **x**_*T*_ from a prior Gaussian distribution, and the reverse diffusion process gradually converts the prior configurations into ensemble configurations **x**_0_. ExEnDiff incorporates a correction term into the pretrained score function **s**_*θ*_(**x**_*t*_,*t*), from measurements **y** from various experimental techniques, including NMR, SAXS, and cryo-EM. (c) A guide-free unified protein ensemble sampler typically fails to sample protein conformation spaces that follow the Boltzmann distribution. In contrast, by guiding the sampling process of a pretrained unified protein ensemble sampler with experimental measurements, ExEnDiff generates configurations that get more closely align with measurement distribution *p*(**y**) or measurement average 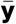 through the reverse diffusion process. This enables ExEnDiff to effectively explore conformation spaces, producing protein conformation ensembles that are closer to the true Boltzmann distribution.

In this work, we employ Str2str^10^ as the pretrained unified protein ensemble sampler due to its high performance in generating feasible protein ensemble configuration. However, it is important to emphasize that ExEnDiff is not confined to a specific unified protein ensemble sampler; it is designed to be adaptable to any ensemble sampler that leverages a diffusion model. As advancements are made in pretrained unified protein ensemble samplers, ExEnDiff will likewise benefit, further enhancing its performance.

### Experimental Data Improves the Accuracy and Diversity of Generated Protein Conformations

To demonstrate the effectiveness of ExEnDiff, we assembled a comprehensive benchmark set of proteins, comprising two ordered proteins (BBA, ^23^ Villin^24^) and eight intrinsically disordered proteins (IDPs), including ACTR,^25^ drkN SH3,^26^ NTail, ^27^ A*β*40,^28^ the ParE2-associated antitoxin PaaA2, ^20^ the intrinsically disordered region from the *Saccharomyces cerevisiae* transcription factor Ash1, ^29^ RS peptide, and ELF3.^30^ This diverse set of proteins contains a diverse range of secondary structures, including *α*−helices and *β*−strands, and various lengths from 24 to 234 amino acids. Furthermore, this benchmark set represents a broad spectrum of proteins, from ordered to highly disordered, and poses a significant challenge for current unified protein ensemble samplers. The ground truth Boltzmann distribution was obtained from long, unbiased MD simulations, except for ELF3, where it was derived using REST (Replica Exchange with Solute Tempering) simulations to enhance sampling of diverse conformational states (simulation details of all proteins are included in the Supplementary Section 1, Fig. S1). In this work we have also introduced a large RS-peptide trajectory (net sampling of 1.76 milliseconds, Fig. S2) for the research community, which was generated using the Folding@home distributed computing system. ^31,32^

We examine the ability of ExEnDiff to generate conformational ensembles that follow Boltzmann distributions by utilizing a wide range of experimentally measurable parameters as guiding metrics. Here, we test the performance of ExEnDiff guided by three sets of guiding metrics: 1) distribution of end-to-end distance (EED), 2) distribution of radius of gyration (Rg), and 3) a combination of distribution of radius of gyration and the secondary structure, percent *α*−helix per residue (Rg+SS). These parameters can be inferred from either a single or a combination of experiments, such as Förster Resonance Energy Transfer (FRET),^33–39^ small-angle X-ray scattering (SAXS),^40–44^ small-angle neutron scattering (SANS),^45–48^ circular dichroism (CD) spectroscopy^49–53^ and Nuclear Magnetic Resonance (NMR) experiments, ^54–56^ for which further details have been provided in Supplementary Section 1. For all proteins in the experiment, we generate 100 distinct protein ensemble conformation samples from the unified protein ensemble samplers. The conformation generation takes less than 300 seconds on a single A100 GPU in all experiments. We note that with additional computational resources, this number can be increased to capture more conformation ensembles.

To provide a clear and straightforward indicator of the performance of ExEnDiff guided by different experiments and the guide-free sampler, we present the accuracy using a single metric — sampler score, as shown in Fig. 2(a). This score is calculated based on the discrepancy between the observables produced by the samplers and those from ground truth MD simulations. For a sampler, a lower sampler score indicates the generated protein conformational ensembles produce close agreement in observables from ground truth MD simulations. The sampler score is calculated based on the discrepancies in end-to-end distance, radius of gyration, solventaccessible surface area, root mean square fluctuation (RMSF), and helix percentage between the sampler and ground truth MD simulations, as detailed in the Methods section and Supplementary Section 5. The results show that ExEnDiff, guided by the experimentally measured parameters, significantly enhances sampling quality. This improvement is particularly notable for disordered proteins, where the original sampler struggles. Notably, ExEnDiff guided by both the radius of gyration and secondary structure (Rg+SS) achieves the best performance, as it integrates both global (radius of gyration) and local (secondary structure) information, greatly enhancing the model’s capacity to explore relevant protein conformation space.

**Figure 2.**
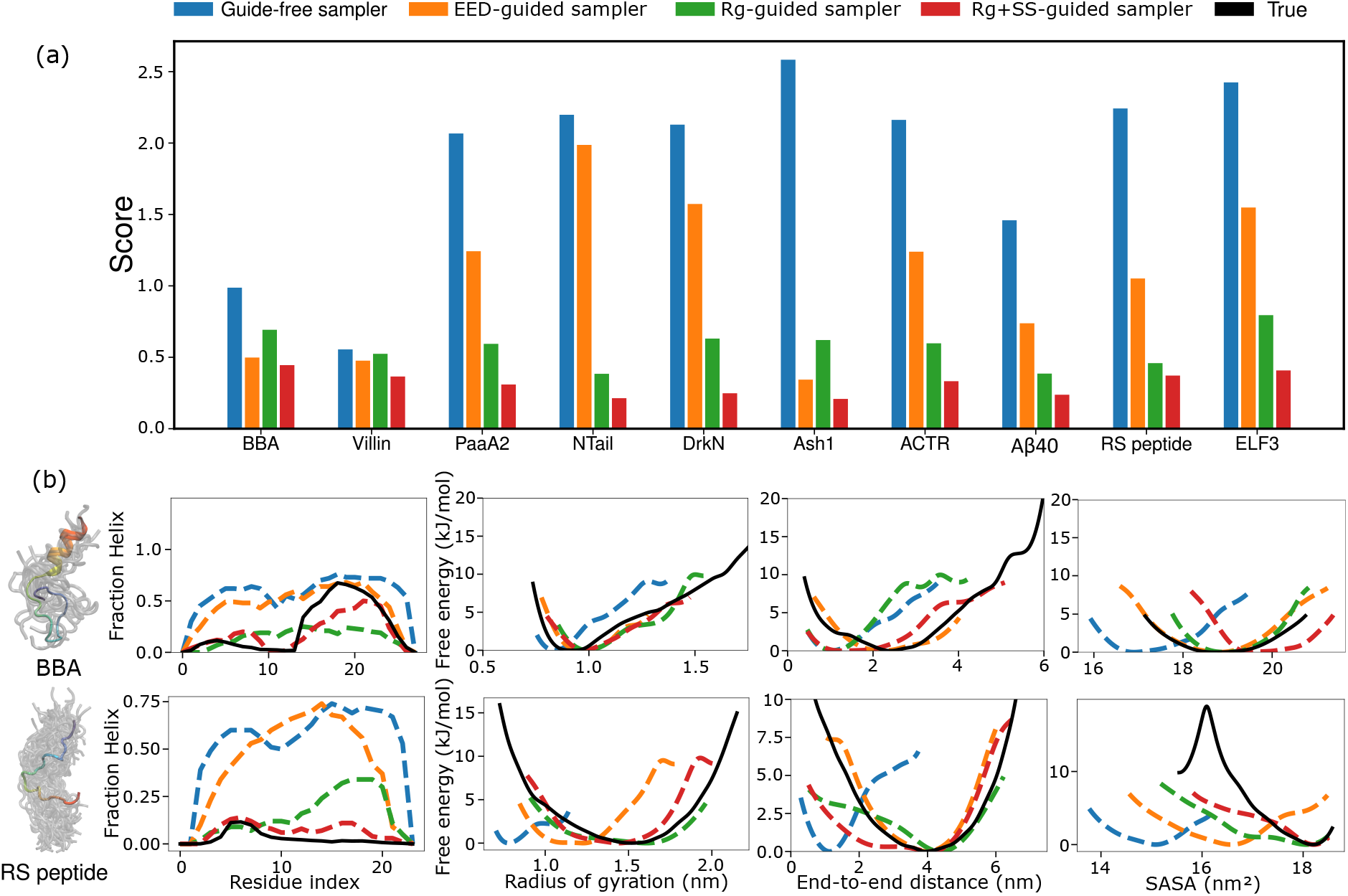
Results for guide-free and different experiment-guided unified protein ensemble sampler EED (end-to-end distance), Rg (radius of gyration), SS (secondary structure)). (a) Unified protein ensemble sampler scores for folded and disordered proteins from the benchmark examined in this work. (b) Fraction helix, FES of radius of gyration, solvent-accessible surface area, and end-to-end distance of two representative proteins from the benchmark: BBA and RS peptide.

We further demonstrate critical physical features using BBA and RS peptide in Fig. 2(b) as representatives of their particularly class of protein: ordered and disordered, respectively (results for all proteins studied are available in Supplementary Section 7.1). While the guide-free sampler performs well in generating feasible ensembles for BBA, ExEnDiff improves the model by refining local secondary structures. Conversely, the guide-free sampler performs poorly on RS peptide, a highly disordered protein, where it tends to sample helix-like structures. This illustrates that a unified protein ensemble sampler trained on PDB data tends to generate fold-like conformations. With the inclusion of experimental measurements, ExEnDiff generates ensembles with more accurate physical features. Importantly, even guiding the sampler with a single parameter, such as the radius of gyration, can significantly improve both global and local aspects of the generated protein ensembles.

We note that all experimental measurements used to guide the protein ensemble generation process in this study are synthetically derived from ground truth MD simulations. In real experiments, secondary structure populations per residue can be accurately predicted from NMR chemical shifts using the *δ*2d program. ^57^ For end-to-end distance and radius of gyration, we provide a preliminary discussion on utilizing real experimental measurements when certain key features are missing in Supplementary Section 6.

### Capturing Mutant Protein Conformations Using ExEnDiff

Generating accurate protein conformational ensembles for protein mutants has long posed a challenge for deep learning models trained on the PDB. Small changes in protein sequences can lead to significant structural discrepancies, which deep learning models can struggle to capture effectively. In this section, we evaluate ExEnDiff’s ability to generate structurally distinct protein conformational ensembles for a mutant of Chignolin. ^58^ This mutant replaces the original tyrosine residues at both termini with glycine and features a threonine-to-proline mutation at position eight, hereon it will be referred to as T8P. These mutations are known to destabilize the native hairpin conformation, ^59,60^ providing a rigorous test of ExEnDiff’s capability in handling structurally divergent mutants.

While the guide-free sampler accurately identifies the native structure of wild-type Chignolin and the local destabilization at the mutation site, it fails to account for the global structural changes induced by the mutation, producing similar conformations for the mutant, as shown in Fig. 3. In contrast, when the sampling process is guided by the radius of gyration, ExEnDiff successfully captures the global destabilization of the *β*-hairpin in the T8P mutant, as reflected in the fraction of *β* structure. Guidance by radius of gyration alone, also improves the prediction of end-to-end distance and solvent-accessible surface area, providing a more accurate representation of the mutant’s conformational ensemble.

**Figure 3.**
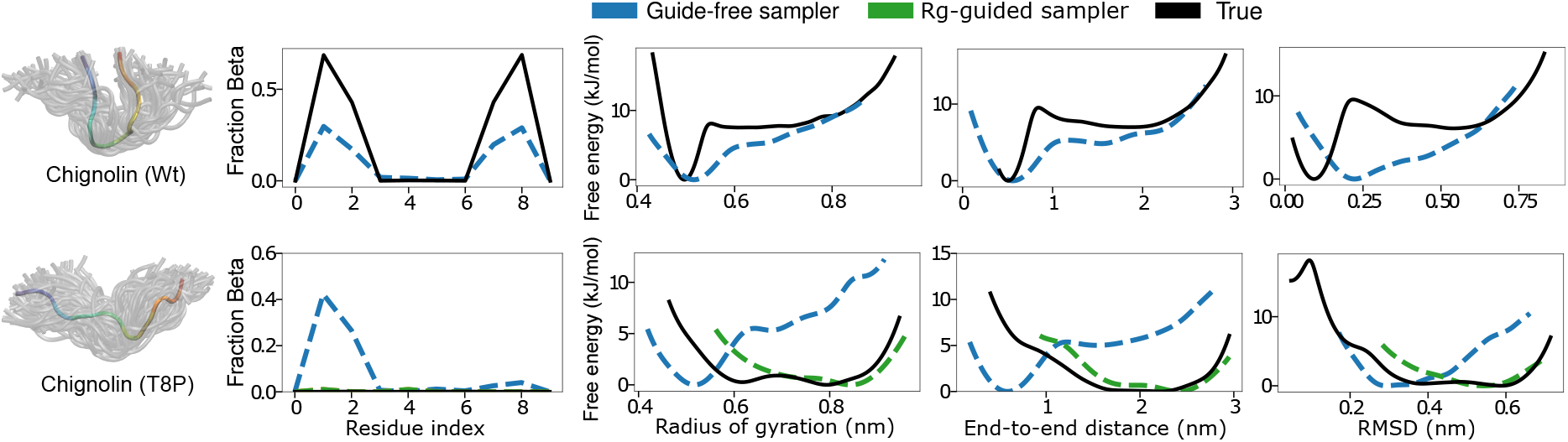
Performance of ExEnDiff guided by radius-of-gyration in capturing mutants’ effect, evaluated by fraction helix, FES of radius of gyration, solvent-accessible surface area, and end-to-end distance using wild type Chignolin and its T8P mutant.

### Cryo-EM 2D Images Enhance the Sampling of Diverse and Physically Realistic Conformations

As a final proof-of-concept of an experiment ExEnDiff can be guided by, we extend the capacity of our model to use 2D images from cryo-electron microscopy (cryo-EM) ^61–66^ as guiding parameters. Cryo-EM has become an important experimental technique for generating high-resolution 3D structures of biological molecules. In cryo-EM experiments, hundreds of thousands of noisy 2D images with random orientations are captured of a sample. Recent studies have suggested that cryo-EM has the potential to extract conformational ensembles that follow the equilibrium distribution of the sample at the temperature before freezing. ^67–73^ These methods either use reconstructed map to guide MD simulations, or combine 2D images and MD simulations as a prior for ensemble reweighting. Either process can become computationally prohibitive when dealing with large-scale proteins.

In this work, we use cryo-EM 2D images as guidance for the unified protein ensemble sampler. We adopt a simple model to represent the configuration-to-image formulation process in cryo-EM following previous work: ^67^

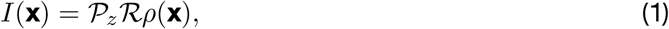

where *ρ*(**x**) is the electron density as a function of the protein configuration **x**, ℛ the rotation matrix of the electron density, and 𝒫_*z*_ the projection along *z* axis. For simplicity, the electron density *ρ*(**x**) is modelled as the sum of spherically symmetric 3D normal densities centered on the *Cα* of the protein.

We generate synthetic images corresponding to each configuration from the ground truth MD simulations, and add multiple levels of noise to form the cryo-EM dataset that guides the sampling process of unified protein ensemble samplers. In this experiment, we assess the performance of ExEnDiff when guided by cryo-EM images across three different signal-to-noise ratio (*snr*) levels: [1, 0.1, ∞], where *snr* = ∞ represents a noise-free scenario. The results, illustrated in Fig. 4 with additional experiments available in the Supplement, reveal several key findings. While the guide-free sampler generates configurations with limited diversity, as evidenced by the radius of gyration, ExEnDiff demonstrates a significantly enhanced ability to explore the protein conformation space, particularly in capturing less compact conformations. Additionally, ExEnDiff shows a strong capacity for predicting correct secondary structure, as indicated by the fraction of helical content.

**Figure 4.**
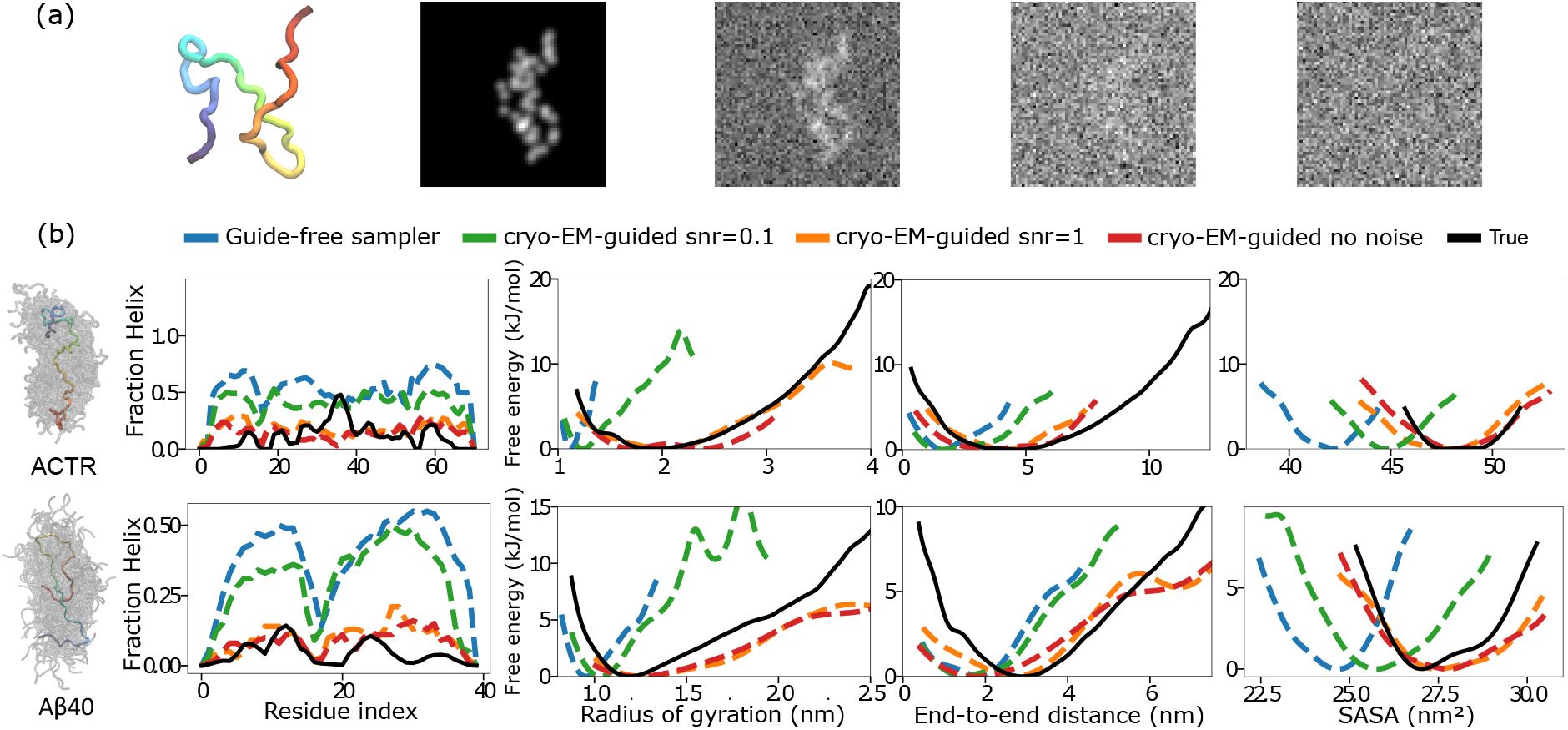
(a) Visualization of synthetic cryo-EM 2D images of an A*β*40 configuration with different levels of signal-to-noise-ratio (*snr*): from left to right is *snr* = ∞ (no noise), *snr* = 1, *snr* = 0.1, *snr* = 0.01. (b) Performance of ExEnDiff guided by cryo-EM 2D density images of *snr* = ∞ (no noise), *snr* = 1, *snr* = 0.1, evaluated by fraction helix, FES of radius of gyration, solvent-accessible surface area, and end-to-end distance of two representative proteins from the benchmark: ACTR and A*β*40.

Remarkably, ExEnDiff produces similar results when guided by cryo-EM images with no noise (*snr* = ∞) and those with an *snr* of 1, indicating its robustness to noisy inputs. This suggests that ExEnDiff has substantial potential for application to real cryo-EM data, which typically contains significant levels of noise. We hypothesize that this resilience to noise arises from the nature of the reverse diffusion process, which inherently functions as a denoising mechanism, thereby enabling ExEnDiff to effectively mitigate noise in cryo-EM images during the protein conformation generation process.

However, we would like to emphasize that experimental guidance using cryo-EM images is still in its initial proof-of-concept stage, and several limitations, further discussed in Supplementary Section 6, must be addressed in future work before this method can be effectively applied to real noisy micrographs.

### Efficient Training of Data-Driven Collective Variables with ExEnDiff

In the final part of the experiment, we showcase a potential application of ExEnDiff, namely the development of efficient data-driven collective variables (CVs) for enhanced sampling simulations. ^74–85^ Compared to traditional physics-based CVs, data-driven CVs can more effectively capture critical low-dimensional features of complex systems. However, a significant challenge with ML-based CVs lies in the quality of the training data, which is often limited by insufficient exploration of the full conformation space of proteins. The capacity of ExEnDiff to generate protein conformational ensembles that reasonably cover protein conformation space makes it an appealing tool as the starting point for training data-driven collective variables for protein ensembles. Inspired by AlphaFold2-RAVE, ^86^ we used ExEnDiff to generate 100 initial samples of a protein conformational ensemble, guided by the radius of gyration and secondary structure. Each sample was subsequently subjected to a short 5 ns MD simulation, resulting in a cumulative 500 ns of MD simulation data. This data was then used to train 2D data-driven CVs. For the CV training, we adopted MESA, ^87^ one of the earliest deep learning approaches that uses an autoencoder-based framework to extract low-dimensional features as CVs. Details of CV training can be found in Supplementary Section 7.3. The same data collection process was applied to the guide-free protein ensemble sampler, and we used the same neural network architecture to train the CVs for both samplers and the converged MD.

One key indicator that a CV is effective is the clear separation of major states into distinct clusters, which reflects the capture of significant configurational similarities and differences. In this study, we applied the k-means clustering algorithm to identify three major conformational states from the converged ground truth MD simulation (a 325 *µ*s MD simulation by DE Shaw Research). These clusters represent distinct secondary structures, with specific helix patterns, as shown in Fig. 5(a). To assess the quality of the CVs, we projected the configurations from the ground truth MD trajectory onto the 2D CV space of each sampler and plotted the corresponding free energy surface (FES), as illustrated in Fig. 5(b). Compared to long unbiased MD simulations, which require microsecond timescales, ExEnDiff identifies multiple major states using significantly fewer computational resources. In contrast, the guide-free sampler captures only a single major conformational state, highlighting its limited ability to detect diverse states.

**Figure 5.**
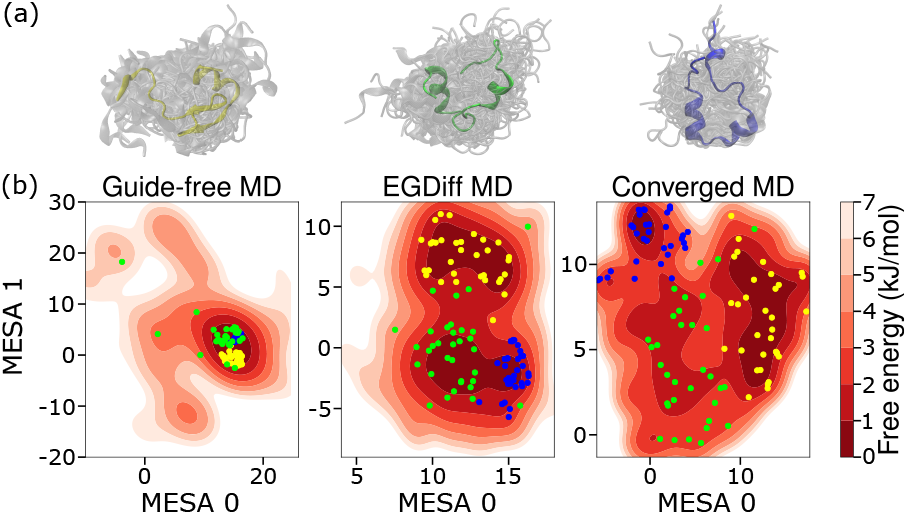
(a) The three major conformational states of Villin, identified by the k-means clustering algorithm, are as follows: the yellow cluster predominantly features ensembles with one major helix at the start of the protein; the green cluster includes ensembles with two major helices, one at the start and one in the middle; and the blue cluster represents ensembles with three major helices located at the start, middle, and end of the protein. (b) Free Energy Surface (FES) of the 2D MESA collective variables (CVs) obtained from training on three independent MD trajectories for Villin. The datasets include: (1) 100 unique starting configurations generated by the guide-free sampler, each run for 5 ns, resulting in a total of 500 ns of MD simulation (guide-free MD); (2) 100 unique starting configurations generated by the ExEnDiff sampler, guided by the radius of gyration and secondary structure, each run for 5 ns, resulting in 500 ns of MD simulation (ExEnDiff MD); (3) a 325 *µ*s MD simulation by DE Shaw Research (Converged MD). The dots, in three different colors, represent the three major conformational states clustered as described in (a), and projected onto each FES.

**Figure 6.**
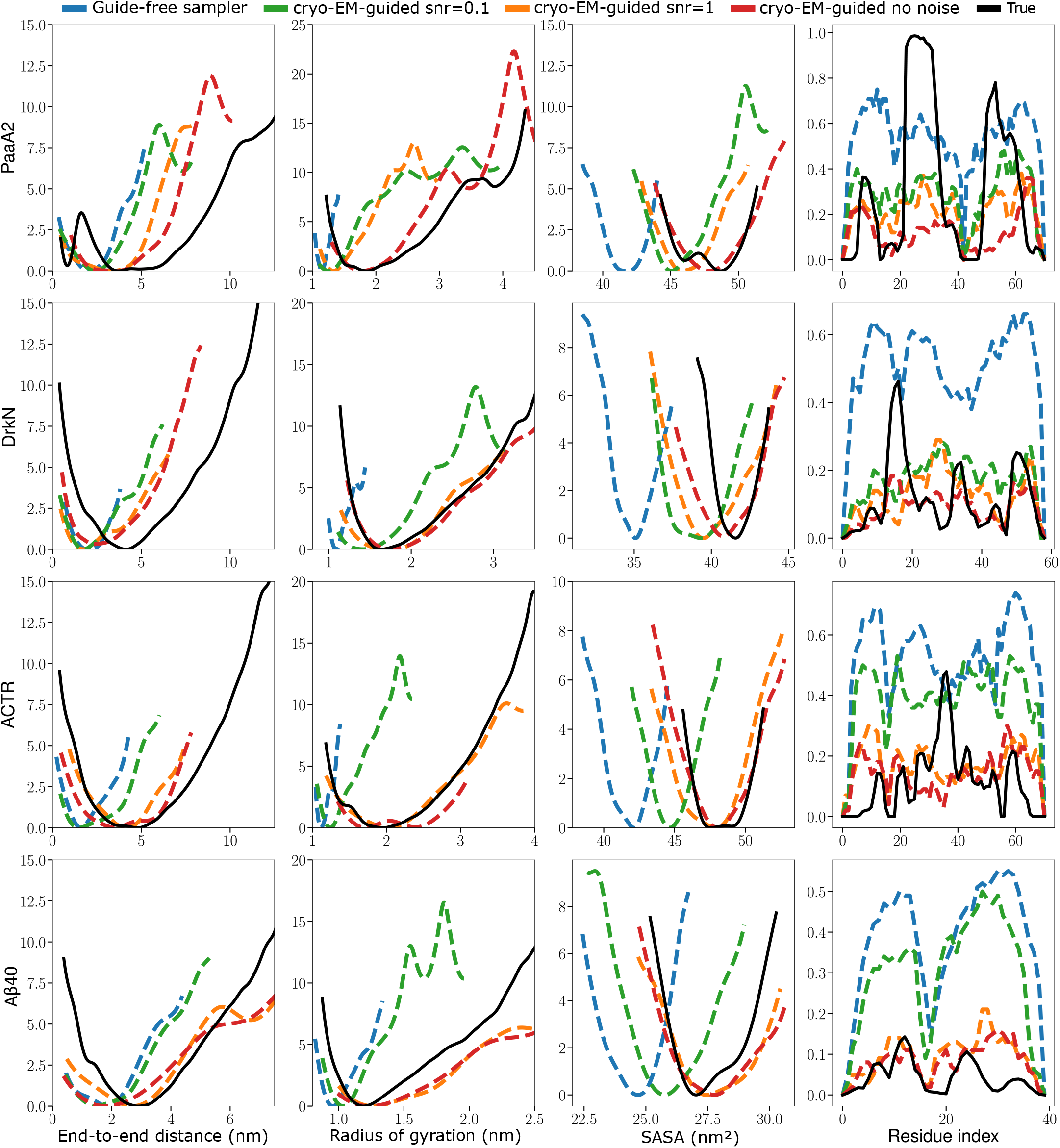
Additional experiment result from protein Ash1, ACTR, A*β*40, RS peptide, and ELF3

**Figure 7.**
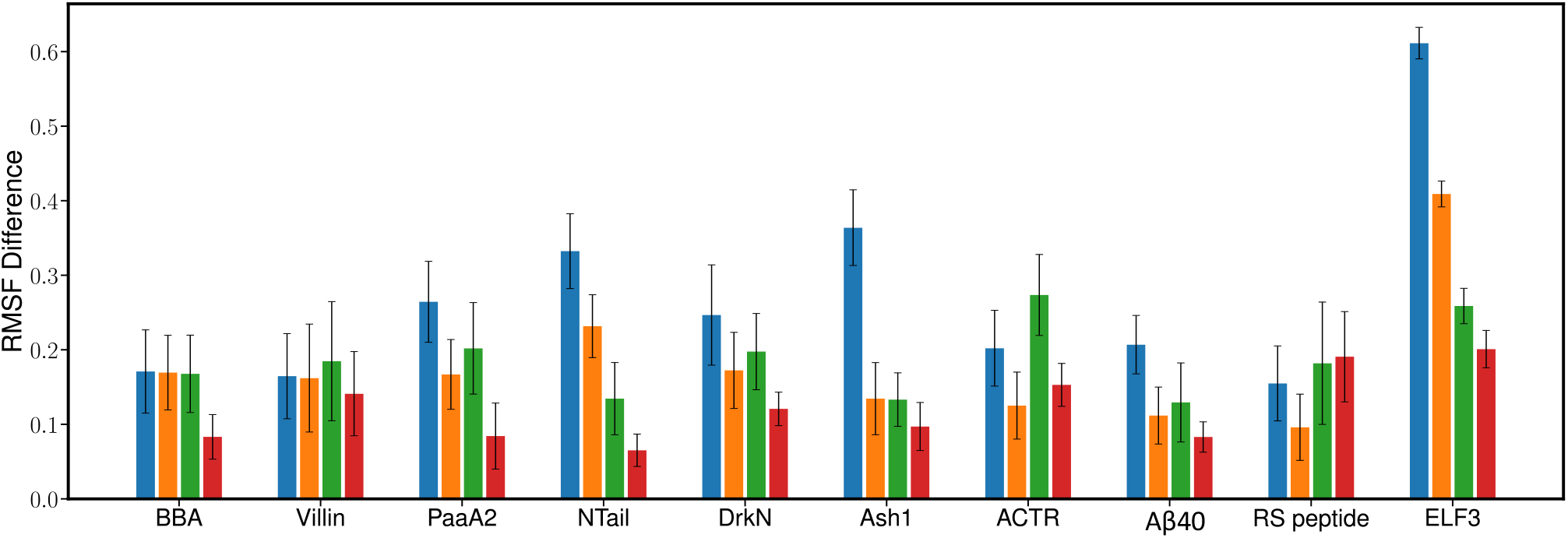
RMSF difference between the ground truth and each sampler: the bar represents the mean RMSF difference and the error bar represents the standard deviation.

**Figure 8.**
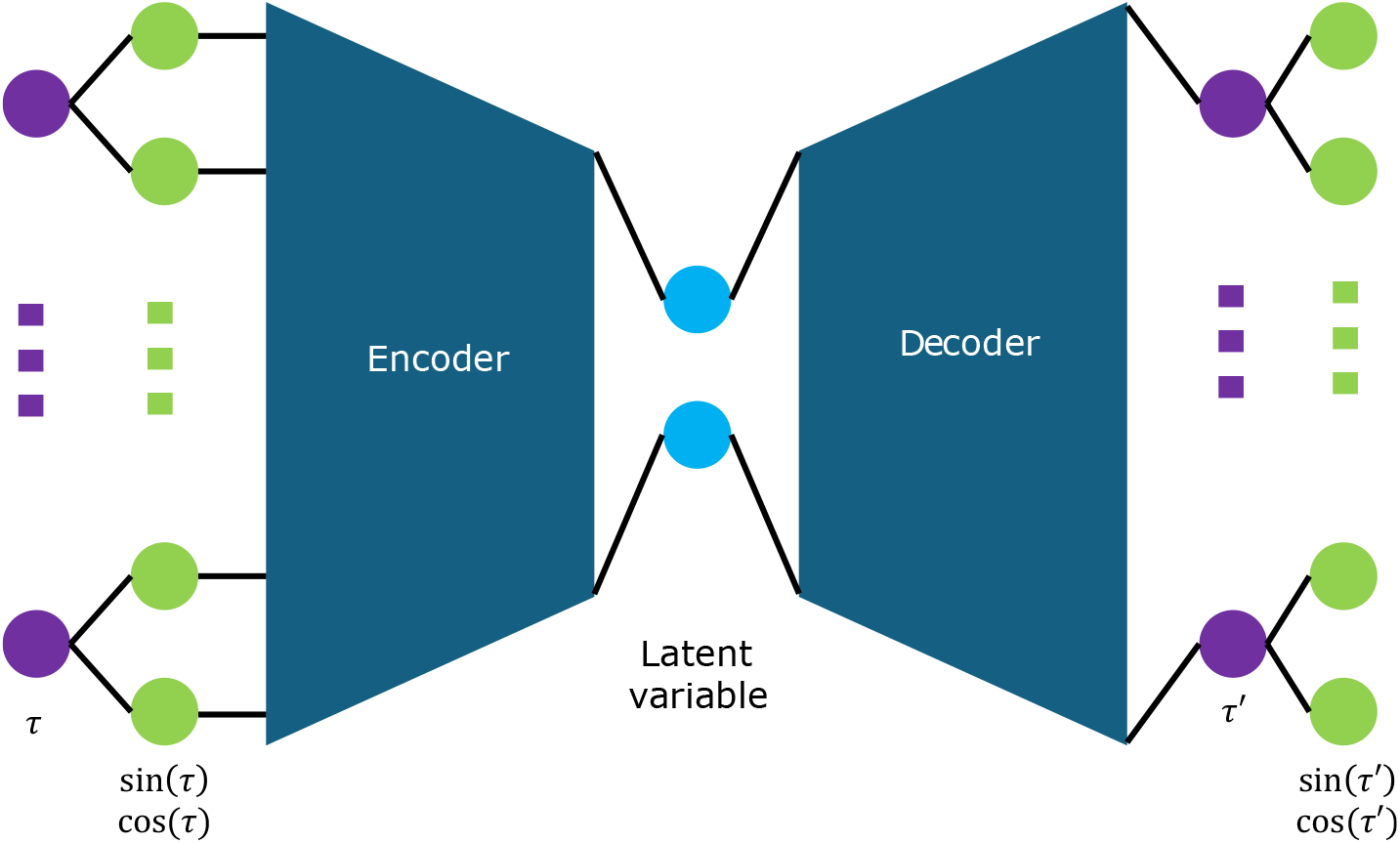
Visual representation of the MESA collective variable (CV) training framework. Input torsion angles, computed from *Cα* atoms, are first embedded using sine and cosine transformations. These embedded values are passed through an encoder to obtain the 2-dimensional latent variables, which are then fed into a decoder to reconstruct the torsion angles. The reconstruction loss is calculated by comparing the recovered sine and cosine transformed torsion angles with the original input angles. The convergedd 2-dimensional latent variables are taken as the collective variables for the system.

**Table 1:** The correction term hyperparameter for end-to-end distance (EED), radius of gyration (Rg), secondary structure (SS), and cryo-EM 2D density images.

|  | EED | Rg | SS | cryo-EM |
| --- | --- | --- | --- | --- |
| $k'_i$ | 5 | 5 | 1 | 500 |

## Discussion and Conclusions

In this work, we have introduced ExEnDiff, a unified protein ensemble sampler that can generate protein ensemble configurations aligning with correct Boltzmann distribution by incorporating experimentally-inferred data. At its core, ExEnDiff’s use of experimental guidance shares significant similarities with molecular dynamics simulations with experimental restraints. In MD simulations, experimental restraints are often used to constrain or guide the conformational sampling through restraining certain variables (like distances, angles, or volumes) to ensure that the simulated ensemble remains consistent with experimental observations. This technique is especially valuable when certain aspects of the system’s conformational space are not well-sampled or when high-resolution data is limited. Experimental restraints in MD can take many forms — distance and angular restraints from chemical shifts, dipolar couplings, and NOE (Nuclear Overhauser Effect) data; cryo-EM or X-ray crystallography derieved restraints aimed to ensure simulated structure fits the density maps and atomic positions; Förster resonance energy transfer (FRET), hydrogen-deuterium exchange, and SAXS (small-angle X-ray scattering) to restrain distances and shapes.^88–92^ Similarly explicit corrective potentials derived from secondary structure assignments have been used to guide MD simulations.^93^ While the experimental guides introduced in our method follow a similar philosophy, the key advantage is a more efficient exploration of conformational space by directly generating diverse ensembles based on experimental data, without running expensive full MD simulations.

We anticipate a broad range of future work building on the ExEnDiff framework. As reported in the Results section, ExEnDiff excels in generating physically meaningful protein conformational ensembles, thereby enhancing the exploration of protein conformation space. This makes ExEnDiff an ideal engine for generating training data for data-driven CVs.

Another significant research direction is the optimization of the force field parameterization pipeline. Traditionally, force field parameterization involves iteratively adjusting force field parameters to align with experimental measurements, followed by rerunning MD simulations using the updated force field. This process is both time-consuming and computationally expensive. Recent advances in deep generative models have shown promise in parameterizing potential energy functions.^94,95^ However, these methods are constrained by their reliance on data from converged MD simulations, limiting their ability to improve beyond the force field that generated the initial MD data. ExEnDiff offers a novel solution by generating protein conformation ensembles that align with various experimental measurements. By integrating ExEnDiff’s ability to produce experimentally consistent and Boltzmann-weighted ensembles with traditional force field components that maintain the underlying physical principles, along with deep generative models that accurately compute full-body potentials, we can achieve a more efficient and accurate approach to protein force field parameterization. This combination could represent a substantial advancement in the field, significantly reducing the time and computational resources required while improving the quality of the resulting force fields.

ExEnDiff can be particularly useful in the data analysis pipelines for single particle cryo-EM, that in principal allows access to the underlying conformational ensemble of the sample making it invaluable for studying proteins and complexes that exist in multiple functional states. A long standing problem in this direction has been to extract free energy profiles from cryo-EM particles.^96^ More recently, an ensemble reweighting approach has been developed that estimates ensemble density from 2D cryo-EM particle data by reweighting based on prior molecular dynamics ensembles.^67^ These approaches rely on a prior, generally from MD, to thoroughly sample the full conformational ensemble, which can be hard for larger proteins and complexes — the target systems in most cryo-EM experiments. Here, ExEnDiff and ExEnDiff supplemented with short MD can provide an attractive alternative by presenting an informative prior through a more efficient exploration of conformational space.

Beyond cryo-EM, along the same vein, the ensembles generated from ExEnDiff can be further refined by integrating a broad range of direct experimental measurments through tools such as HDXer and BioEN.^97–101^

In the current version, ExEnDiff focused on structural information obtained after processing raw experimental data, often through established physical and mathematical models. For example, a commonly used method to extract Rg from SAXS data is the slope from the Guinier plot at small scattering vector regimes.^102–104^ However, this approach comes with several assumptions, such as the homogeneity and compactness of the sample. For complex or irregularly shaped particles (e.g., elongated, branched, or highly anisotropic structures), the Guinier approximation may not be applicable, leading to inaccuracies in the structural interpretation. Therefore, arguably, ExEnDiff with raw experimental data, rather than processed outputs, could be more informative at guiding towards relevant protein structures. Given that ExEnDiff is a versatile tool capable of handling diverse data types, as previously demonstrated, future iterations could take advantage of this flexibility by directly incorporating raw experimental data into the inference process. This would allow ExEnDiff to overcome the limitations of or supplement traditional model-based methods.

Despite its significantly improved performance, ExEnDiff has a few limitations. One is the use of a simple harmonic restraint potential in manifold constraint sampling, which ignores covariance between dimensions. A more advanced approach, such as Mahalanobis-distance-based restraints, could address this problem and will be explored in future iterations. Another limitation is that ExEnDiff relies on samples from the measurement distribution rather than the full distribution itself, which can be problematic for high-dimensional data such as the 2D density images obtained from cryo-EM experiments, if the samples are not fully representative. To mitigate this, future efforts will focus on developing a density-guided diffusion model. In this vein, another promising future direction is to explore the model’s limits with larger protein structures, and extending its capability to handle polynucleotides and biologically relevant macromolecular assemblies.

Overall, we have demonstrated that ExEnDiff is an effective tool for accessing conformational distributions of protein ensembles using simple experimentally-obtained values as a guide. We believe this approach will be key to pushing the study of protein ensembles into the data regime required for the large-scale learning of not just static structures but true protein ensembles. This will accelerate our ability to understand the effects of protein environment and mutations on protein ensemble properties, currently an outstanding problem in the field, with tremendous implications for both basic science and human health.

## Materials and Methods

### Score-based diffusion models for unified protein ensemble sampler

The current unified protein ensemble samplers^10–13^ all use diffusion models as the base generative model. The score-based diffusion model perturbs data with original distribution *p*_0_(**x**) to Gaussian distribution *p*_*T*_ (**x**) with a diffusion process over a unit time horizon by a stochastic differential equation (SDE):

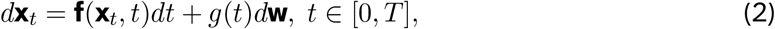

where **f**(**x**_*t*_, *t*), *g*(*t*) are chosen diffusion and drift functions and **w** denotes a standard Wiener process. For any diffusion process in Eq. 2, it has a corresponding reverse-time SDE: ^105^

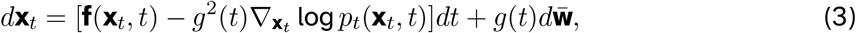

with 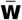 a standard Wiener process in the reverse-time. The score-based diffusion model learns 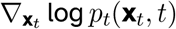 with a parameterized score function **s**_*θ*_(**x**_*t*_, *t*). To sample from the data distribution *p*_0_(**x**), we draw samples from the prior distribution *p*_*T*_ (**x**) ∼ 𝒩 (**0, *I***), and then discretize and solve the reverse-time SDE with numerical methods, e.g. Euler-Maruyama discretization. Details of the score-based diffusion model used in this paper can be found in supplementary Section 2.

The primary objective of all unified protein ensemble samplers is to learn and generate independent and identically distributed (i.i.d.) protein configuration samples from the underlying distribution *p*(**x**|seq), which represents the conformational space associated with a given protein sequence. Achieving this objective is particularly challenging due to the inherent difficulty in obtaining a comprehensive set of proteins with conformation samples that accurately follow the Boltzmann distribution.

### ExEnDiff: A manifold constraint sampling protocol for guiding unified protein ensemble sampler with experimental measurements

In this section we propose ExEnDiff, a simple framework that aims to enhance existing diffusion-based unified protein ensemble sampler’s to generate physically feasible protein conformation samples that follow a true Boltzmann distribution. Built upon the manifold constraint sampling technique, ^106^ a novel experiment-guided sampling strategy is proposed to alter the intermediate score function to match the experimental measurements.

Consider an experimental measurement that can be written as a function of atomic positions: **y** = *A*(**x**_0_), where **y** = [*y*_1_, …, *y*_*m*_] = [*A*_1_(**x**_0_), …, *A*_*m*_(**x**_0_)] represents *m* one dimensional features. With a prior diffusion-based protein ensemble sampler, our aim is to retrieve and generate conformation ensembles from the conditional probability distribution *p*(**x**_0_|*A*(**x**_**0**_) = *y*). If we take the score-based diffusion model as the prior, we can use the reverse diffusion from Eq. 3 as the sampler from the conditional distribution as follows:

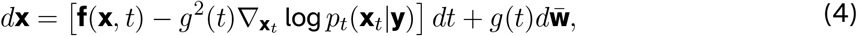

where 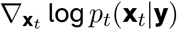 can be decomposed using the Bayes’ rule:

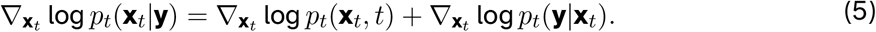

Applying the manifold constraint sampling technique, the posterior term can be estimated with a harmonic restraint potential function:

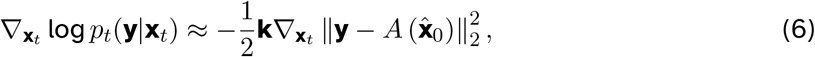

where **k** = [*k*_1_, …, *k*_*i*_], with *k*_*i*_ the restraint constant of *i*−th function, and 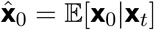 is the posterior mean. Finally, with a pretrained diffusion model **s**_*θ*_(**x**_*t*_, *t*) as the prior, we can approximate 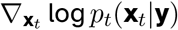 with a conditional score function **s**_rest_(**x**_*t*_, *t*) at the inference time:

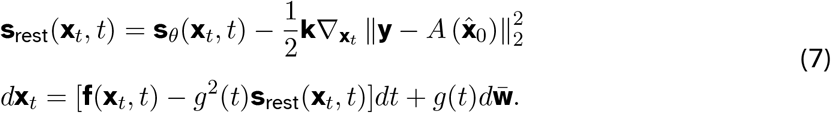

The experimental measurement can give either the probability distribution *P*_measurement_(**y**), or the ensemble average 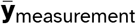. If the experimental measurements give observable average 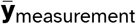, we approximate the probability distribution of **y** with a Gaussian distribution centered on 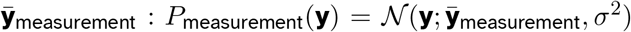, with *σ* a manually-designed variable variance. We want to enhance the protein ensemble sampler’s capacity by forcing it to generate samples that match with the experimental measurement *P*_sample_(**y**) ≈ *P*_measurement_(**y**), where

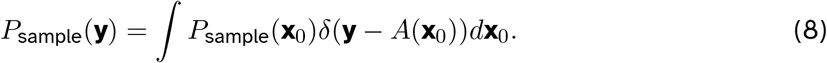

This can be achieved by the following procedure:

- Sample **y**_1_, **y**_2_, …, **y**_*N*_ ∼ *P*_measurement_(**y**).
- For each **y**_*i*_, sample 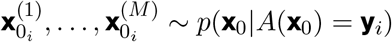.

The first step can be achieved easily, since **y** is usually of low dimension. For the more challenging second step, we utilize a state-of-the-art unified protein ensemble sampler as the prior, and apply manifold constraint sampling techniques. This two-step sampling procedure ensures we have a information-rich protein ensemble sampler to start with, and can greatly enhance the model’s capacity to generate protein ensembles that match with experiment measurements at no additional computational cost. We include full details of ExEnDiff in supplementary Section 3, and choice of restraint constant *k*_*i*_ in supplementary Section 7.2.

### Calculation of sampler score

To compare the relative accuracy of each sampler, we report the sampler scores. The sampler scores for IDP are computed as the following:

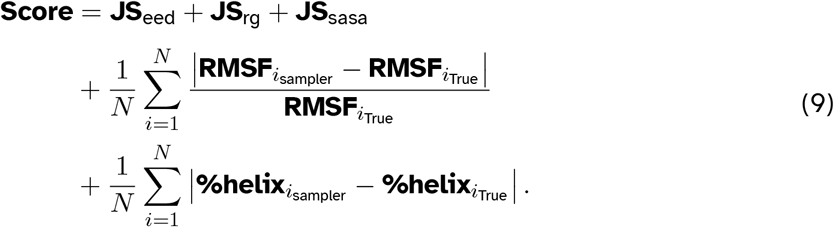

The score is a sum of five terms: the first three terms are the Jensen-Shannon (JS) distances w.r.t the end to end distance, radius of gyration, and the solvent accessible surface area. These JS distances are measured to quantify the similarity between the distributions of protein conformation ensembles generated by the sampler and those of the ground truth. These terms provide a global measure of how well the sampler captures the overall structural features of the protein ensemble. The fourth term is the residue-specific deviation in root-mean-square fluctuation (RMSF), which measures the difference in flexibility at each residue between the sampled and true ensembles. The fifth term captures the discrepancy in helix content per residue, indicating how well the sampler reproduces the secondary structure elements. These latter two metrics focus on local structural differences, providing a finer resolution of the sampler’s accuracy at the residue level. Together, these components offer a comprehensive evaluation of the sampler’s performance in capturing both global and local features of protein conformations.

## Data Availability

The benchmark protein dataset except RS-peptide has been taken from previous published research. RS-peptide trajectory is available for download https://doi.org/10.5281/zenodo.14013285. All the scripts and codes are available https://github.com/flatironinstitute/ExEnDiff

## Acknowledgement

G. Lin would like to thank the support by National Science Foundation (DMS-2053746, DMS-2134209, ECCS-2328241, CBET-2347401 and OAC-2311848), and U.S. Department of Energy (DOE) Office of Science Advanced Scientific Computing Research program DE-SC0023161, and DOE–Fusion Energy Science, under grant number: DE-SC0024583. M.C. acknowledges support from support from the National Science Foundation (EAR-2246687). We would also like to thank the Folding@home community and Géraud Krawezik (software engineer at Flatiron Institute) for helping us set up Folding@home simulations. The Flatiron Institute is a division of the Simons Foundation.

---

**Figure 1.**
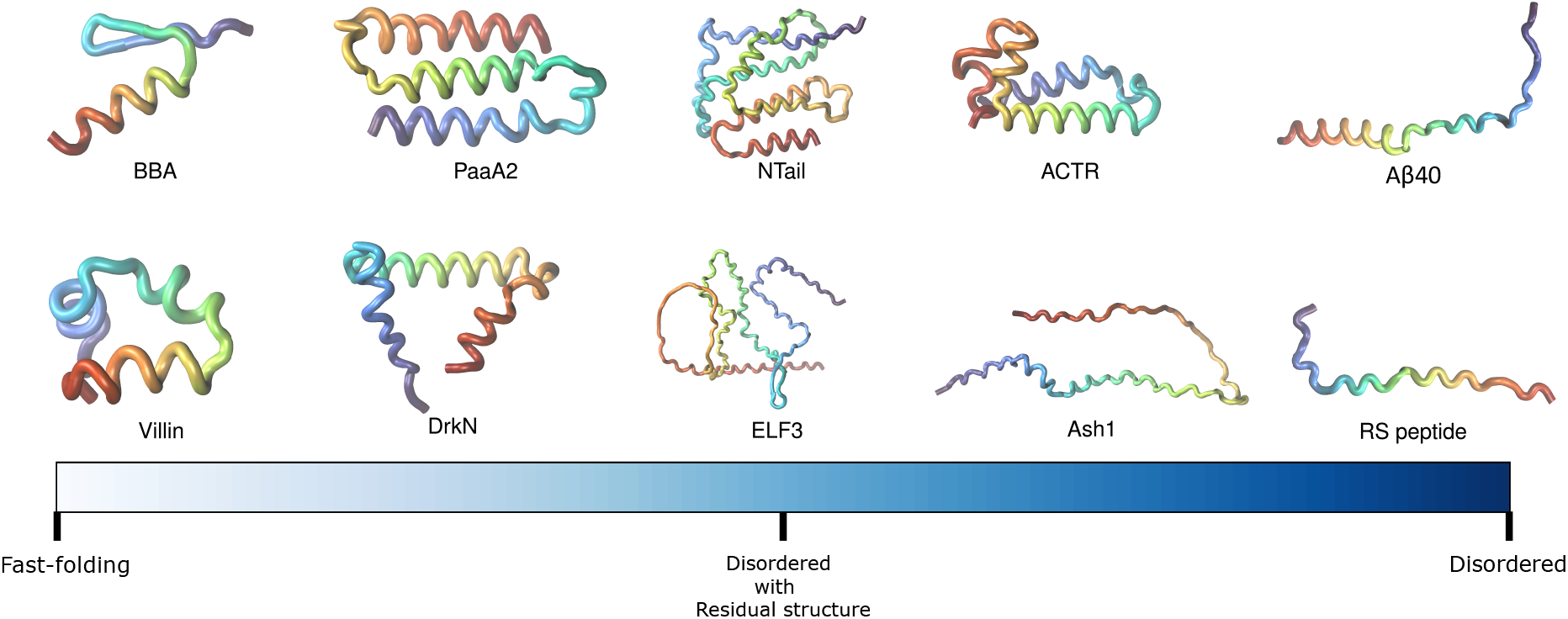
Schematic illustration of benchmark set of proteins used in this work to assess the performance of ExEnDiff.

**Figure 2.**
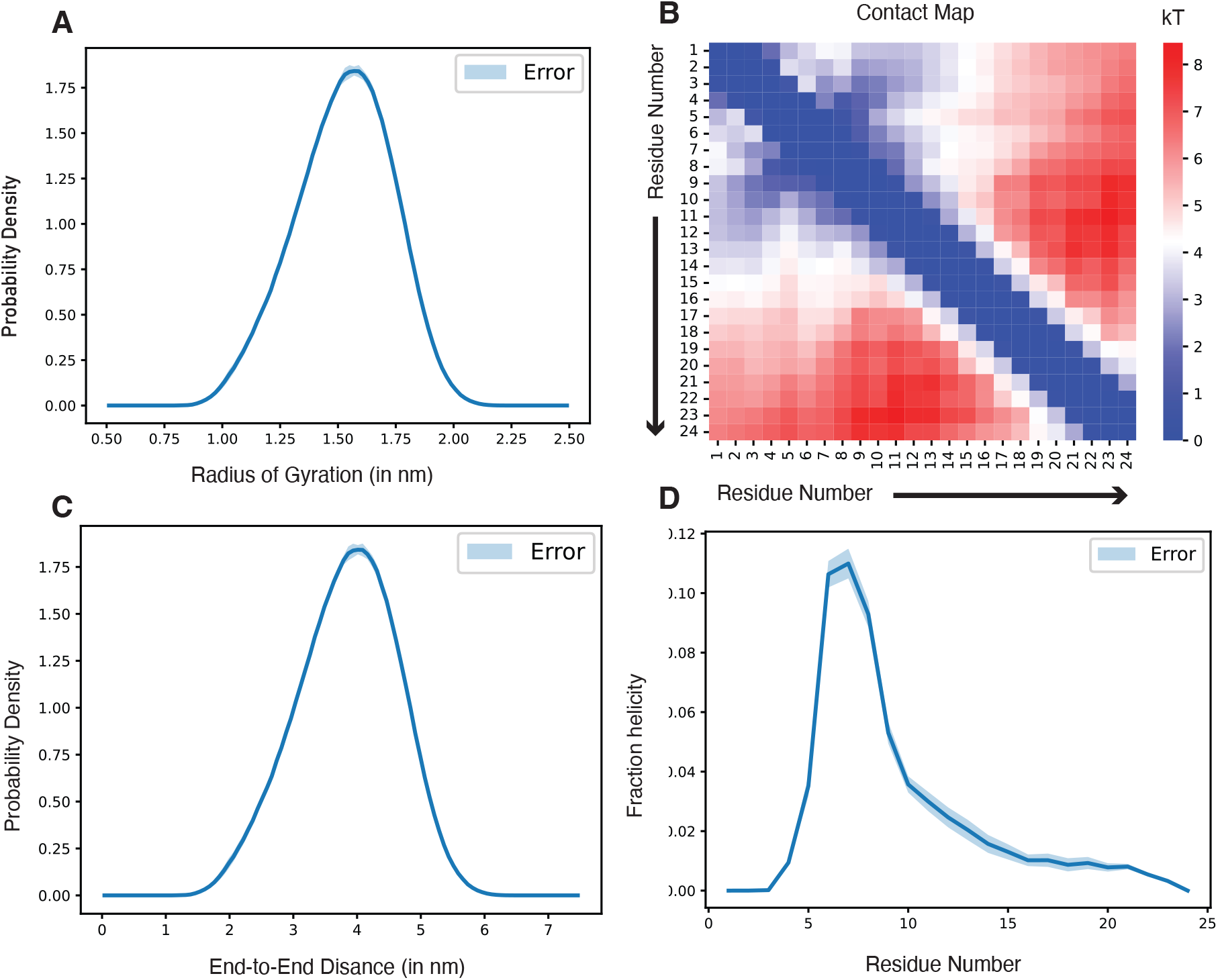
Overview of RS-peptide simulations. **a)** Radius of gyration, **b)** Contact map, **c)** End-to-end distance, **d)** Helical fraction

**Figure 3.**
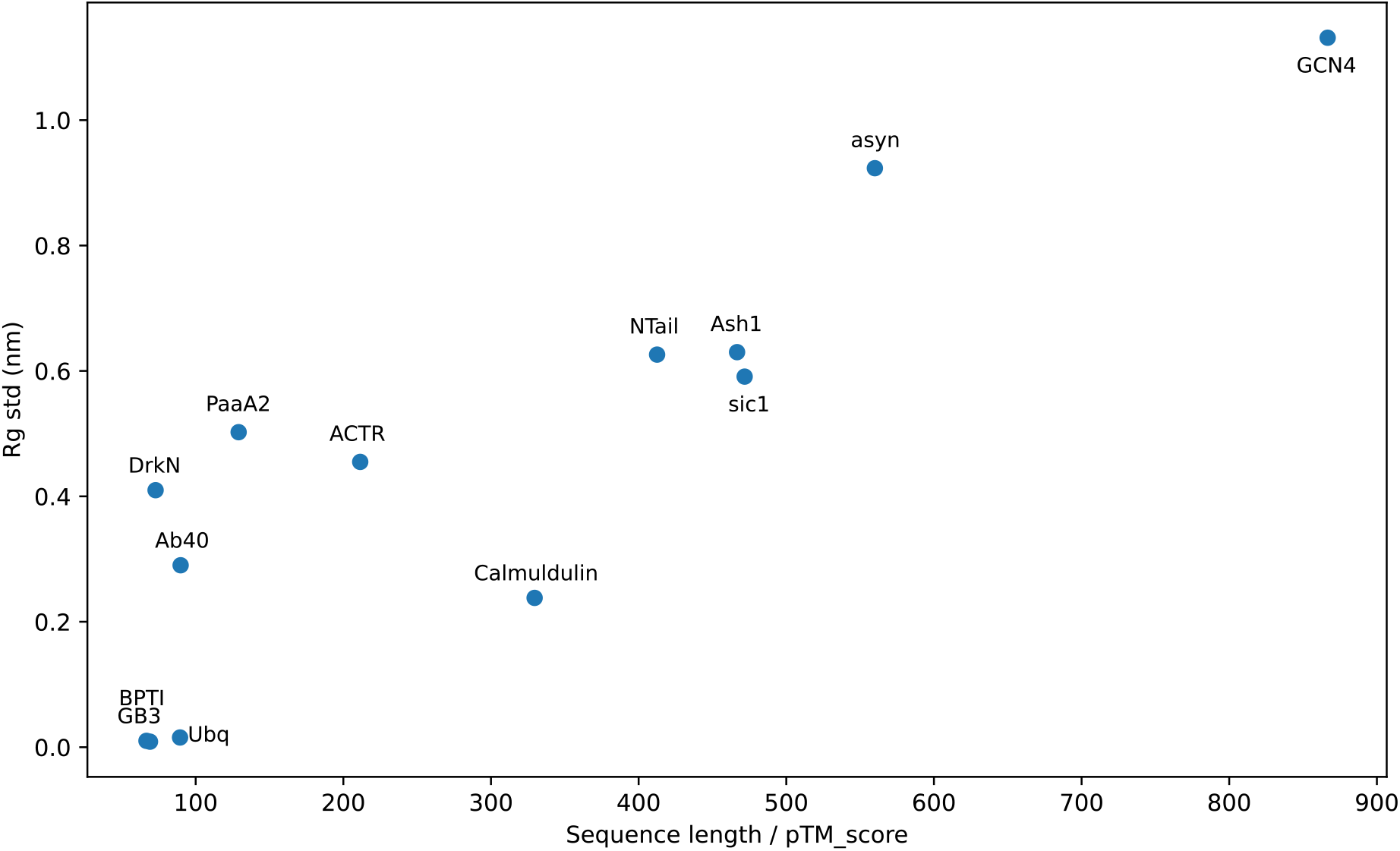
Preliminary investigation of relationship between the radius of gyration (Rg) standard deviation, protein sequence length, and pTM score predicted using Alphafold3.

**Figure 5.**
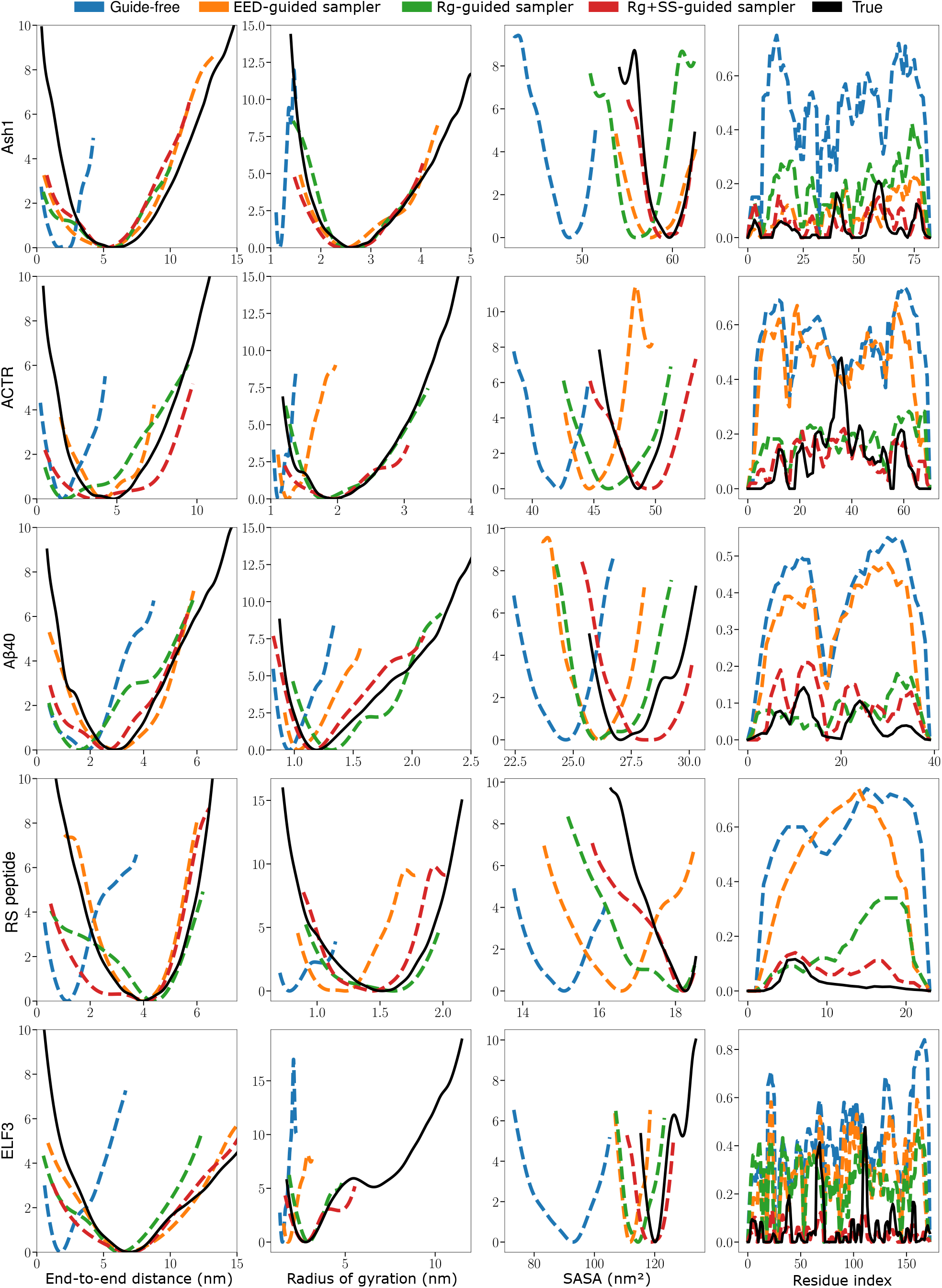
Additional experiment results from protein Ash1, ACTR, A*β*40, RS peptide, and ELF3

## Supporting information

### 1 Benchmark Protein Dataset

We assemble a testing dataset of 10 proteins, from fast-folding proteins to intrinsically disordered proteins, as shown in Fig. 1. The ground truth Boltzmann distribution was obtained from long unbiased MD simulations. Simulations of Villin, ACTR, drkN SH3, NTail, *Aβ*40, PaaA2, and Ash1 were conducted by DE Shaw research using the a99SB-disp force field^107^ for 30 *µs*. The force field parameters were optimized based on experimental measurements to achieve high accuracy in simulation results across various ordered and disordered proteins. Simulations of BBA and Villin were conducted by DE Shaw research^108,109^ using the CHARMM22* force field and the modified TIP3P water model^110^ compatible with the CHARMM^111^ force field for 325 *µs* and 125 *µs* respectively. For ELF3-PrD, the converged free-energy surface was inferred from the REST2 (Replica Exchange with Solute Tempering) simulations, ^112^ which is an unbiased enhanced simulation method. The details of the simulation protocol are listed in Lindsay *et al*. ^30,113^

Finally, for the RS-peptide, we ran long unbiased simulations of up to 2 milliseconds of simulation time, with 20 unique starting structures, each run with 100 clones of ∼0.9 micro-second each using Amber03ws force field^114^ and Tip4p/2005 water model. ^115^ The system was initially solvated, and neutralizing ions were added. Energy minimization was performed using the steepest descent algorithm to remove steric clashes, with a maximum force tolerance of 1000 kJ/mol/nm. The system underwent equilibration in two stages: first, a 100 ps NVT equilibration at 300K, with temperature kept constant using the v-rescale thermostat (with a time constant of 0.1 ps), followed by a 1 ns NPT equilibration at 1 bar with the Parrinello-Rahman barostat (with a time constant of 2.0 ps). The production simulation was run with a 2 fs time step. Electrostatic interactions were treated using Particle Mesh Ewald (PME) with a cutoff of 1.0 nm, and other non-bonded interactions were modified with the Potential-shift-Verlet scheme with a cutoff of 1.2 nm. Constraints on bonds involving hydrogen were applied using the LINCS algorithm. ^116^ The simulations were performed in the Folding@Home distributed computing environment. ^31,32^

Here, we present some tests for convergence and basic structural features from our simulations. The simulations were divided into 20 independent fragments with 86-89 *µs* each, and a distribution of radius-of-gyration, end-to-end distances and helical-fraction per residue was calculated. From Fig. 2 a,c and d, we could note that the peptide exhibits stable convergence across all structural metrics.

In order to demonstrate the applicability of ExEnDiff in studies of point mutation, we used the fast folding protein Chignolin, and its point mutant Chignolin T8P. The simulation of wild-type Chignolin was conducted by DE Shaw research^108,109^ using the CHARMM22* force field and the modified TIP3P water model^110^ compatible with the CHARMM^111^ force field for 106 *µs*. For the Chignolin T8P mutant, we ran a 1.5 *µs* simulation using the a99SB-*disp* forcefield developed by DE-Shaw Research for accurate simulations of both folded and disordered protein, with TIP4P-D water. The atomic structure for the mutant was prepared using Pymol by replacing the Threonine at 8th residue to Proline in the native structure of Chignolin. Following solvation, neutralizing ions were introduced to balance the system’s charge. The simulation protocol is exactly equivalent to the case of RS-peptide, as detailed above.

### 2 Additional details of ExEnDiff

In ExEnDiff, we consider the variance preserving (VP) form of the SDE in Denoising Diffusion Probabilistic Model (DDPM): ^9^

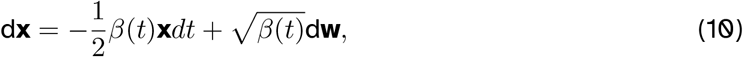

with *β*(*t*) representing the variance schedule. In a discretized setting of DDPM, we define *β*_1_, *β*_2_, …, *β*_*T*_ as the sequence of fixed variance schedule. If we define *α*_*t*_ = 1 − *β*_*t*_ and 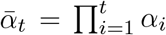, then the forward diffusion process can be written as:

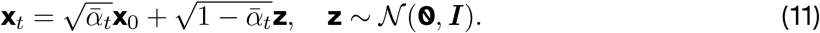

The forward diffusion process with *T* iterations of a DDPM model is defined as a fixed posterior distribution *p*(**x**_1:*T*_ |**x**_0_). Given a list of fixed variance schedule *β*_1_, …, *β*_*T*_, we can define a Markov chain process:

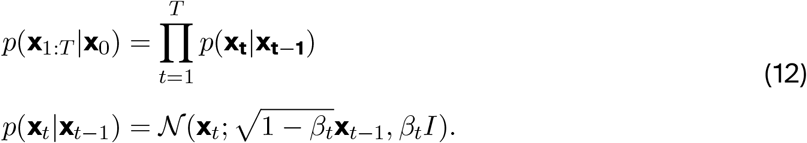

We have the following property:

**Property 1** *The marginal distribution of the forward diffusion process p*(**x**_*t*_|**x**_0_) *can be written as:*

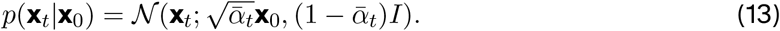

This can be obtained by the following proof:

**Proof 1** *Using p*(**x**_*t*_|**x**_*t*−1_) *from* (12), *we can obtain:*

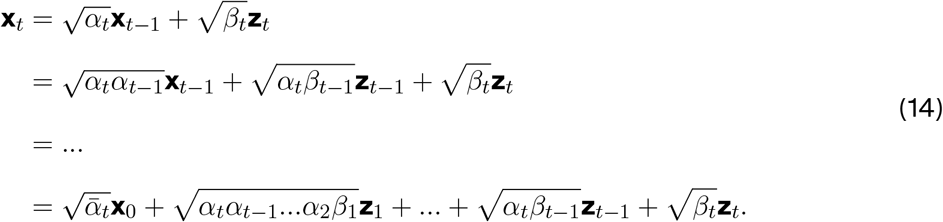

*We can see that p*(**x**_*t*_|**x**_0_) *can be written as a Gaussian with mean* 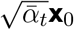 *and variance* 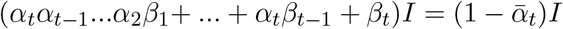.

This property allows us to write the forward diffusion process in the form of Eq. (11). As *T* → ∞, the discretized Eq. (11) converges to the SDE form defined in Eq. (10).

ExEnDiff requires estimations of the posterior mean configuration 𝔼 [**x**_0_ | **x**_*t*_]. This can be obtained using Tweedie’s formula.

**Lemma 1** *(Tweedie’s formula) Let µ be sampled from a prior probability distribution G*(*µ*) *and z* ∼ 𝒩 (*µ, σ*^2^), *the posterior expectation of µ given z is as:*

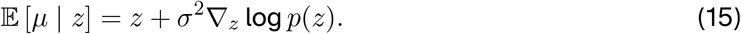

From Tweedie’s formula, we can obtain the following property:

**Property 2** *For DDPM with the marginal distribution p*(**x**_*t*_|**x**_0_) *of the forward diffusion process computed in equation 13, p*(**x**_0_|**x**_*t*_) *has a posterior mean at:*

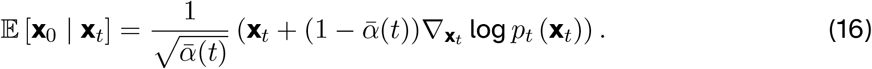

### 3 Restraint potential in molecular dynamics simulation

Consider a mapping function on atomistic coordinates of a protein configuration **x**: **y** = *A*(**x**), where **y** = [*y*_1_, …, *y*_*m*_] = [*A*_1_(**x**), …, *A*_*m*_(**x**)] represents *m* low dimensional features. In molecular dynamics (MD) simulations, generating protein ensembles that correspond to a specific data ensemble value *A*(**x**) = **y** can be achieved by adding a harmonic restraint potential centered on **y**. This restraint potential can be expressed as:

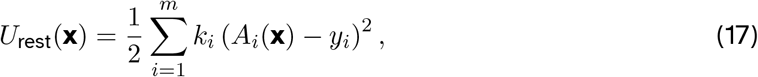

where *k* is the force constant that determines the strength of the restraint, *A*_*i*_(**x**) are the low-dimensional features as functions of the coordinates **x**, and *y*_*i*_ are the target values for these features. The total potential energy of the system, including the restraint, is then given by:

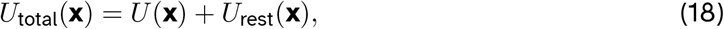

where *U* (**x**) is the original potential energy of the system. By including this restraint potential in the MD simulations, the system is encouraged to explore conformations where the features **y** = *A*(**x**) are close to the desired target values, thereby generating ensembles that satisfy the specific data ensemble constraint *A*(**x**) = **y**.

### 4 Guiding Experiment Parameters

#### End-to-end distance

The end-to-end distance is a crucial parameter in the study of protein structure and dynamics, representing the distance between the N-terminal and C-terminal ends of a protein. This parameter provides insights into the overall conformation and folding state of the protein. Experimentally, the end-to-end distance can be measured using techniques such as Förster Resonance Energy Transfer (FRET),^33–39^ Small-Angle X-ray Scattering (SAXS)^40–44^ and Electron Paramagnetic Resonance (EPR). In FRET, fluorescent dyes are attached to the ends of the protein, and the efficiency of energy transfer between the dyes, which depends on the distance between them, is measured. In SAXS, the scattering pattern of X-rays by the protein in solution is analyzed to obtain information about its size and shape, which through appropriate mathematical and computational models can yield the end-to-end distance. Finally with EPR, spin labels can be covalently attached to the termini, and interaction between them can be used to infer the end-to-end distance.

The end-to-end distance *R* is mathematically defined as:

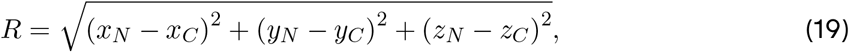

where (*x*_*N*_, *y*_*N*_, *z*_*N*_) and (*x*_*C*_, *y*_*C*_, *z*_*C*_) are the coordinates of the C*α* atoms at the N-terminal and C-terminal ends of the protein, respectively. Understanding the end-to-end distance helps in characterizing the protein’s flexibility, folding pathways, and functional states.

#### Radius of gyration

The radius of gyration (*R*_*g*_) is a critical parameter in the study of protein structure, representing a measure of the protein’s compactness. It is defined as the root-mean-square distance of the atoms from their common center of mass, providing insights into the overall shape and distribution of mass within the protein molecule. The radius of gyration is particularly useful in comparing the compactness of different proteins or the same protein under various conditions, such as different folding states or environments. The *R*_*g*_ can be experimentally measured using techniques like small-angle X-ray scattering (SAXS) or small-angle neutron scattering (SANS).^45–48^ Both these methods involve irradiating the samples with X-ray or neutrons and analyzing scattering pattern at small angles. Then the *R*_*g*_ is determined from the slope at the linear regime in the Guinier plot.

The equation for the radius of gyration is given by:

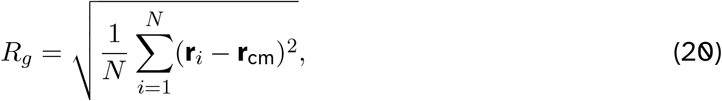

where *N* is the number of atoms, **r**_*i*_ is the position vector of the *i*-th atom, and **r**_cm_ is the position vector of the center of mass of the protein. This equation effectively quantifies the spread of the atomic positions around the center of mass, making the radius of gyration a valuable descriptor in the field of structural biology. In this work, we consider radius-of-gyration only w.r.t the *Cα* atoms.

#### Secondary structure

We use the helix percentage per residue as the guiding parameter. Helix structures in proteins are characterized by a repeating pattern of hydrogen bonds and specific geometric arrangements of amino acids. Experimentally, secondary structure can be calculated using methods such as circular dichroism (CD) spectroscopy^49–53^ and Nuclear magnetic resonance experiments. ^54–56^ Additionally, computational tools such as *δ*2d program^57^ have demonstrated high accuracy predicting secondary structure populations per from NMR chemical shifts measurements.

Let *r* = [*r*_1_, …, *r*_*N*_] denote the secondary structure of an *N* -residue protein, where *P* (*r*_*i*_ = *H*) represents the probability that the *i*-th residue is in a helical state. We divide the sequence into *m* “helix groups” and denote the index of the residue with the highest helix probability within each group as *G*_*i*_. For each sample, the residue corresponding to *G*_*i*_ is sampled first according to its distribution 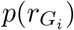 Within each helix group, the probability that a residue is part of a helix is determined sequentially. For instance, for the residue at position *G*_*i*_ − 1, the probability of being in a helical state given that the residue at position *G*_*i*_ is in a helical state, 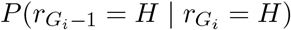, is calculated as follows:

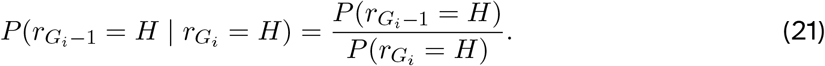

Similarly, we can write 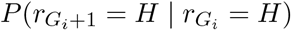 as follows:

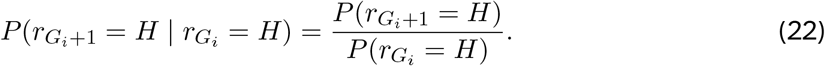

This sequential approach ensures that the helical state of residues within each group is influenced by the residue with the highest helix probability, allowing for the propagation of helical propensity throughout the group in an orderly manner, thus avoiding unrealistic helix assignment.

One common type of helical structure is the alpha helix, where the i-th residue forms a hydrogen bond with the (i+4)-th residue. This creates a compact, right-handed helical structure with 3.6 residues per turn. The distance between the i-th and (i+3.6)-th residues in an alpha helix is typically fixed, reflecting the regularity of the helical backbone. This fixed distance is approximately 5.4 Å, which corresponds to the pitch of the helix—the vertical rise per complete turn of the helix. This consistent spacing is crucial for the stability and regularity of the helix, contributing to the overall structural integrity and functional properties of the protein. The precise distance can be slightly altered depending on the specific sequence and environmental factors, but the helical geometry is generally preserved across a wide range of protein structures. To induce helix formation at the *i*-th residue, we set the *Cα* − *Cα* distance between the *i* and *i* + 4 residues to 6.3 Å, as described by the following equation:

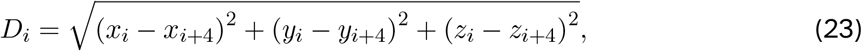

where *D*_*i*_ represents the distance between the *i*-th and *i* + 4-th residues. To enforce this helical structure, we apply an adapted manifold constraint:

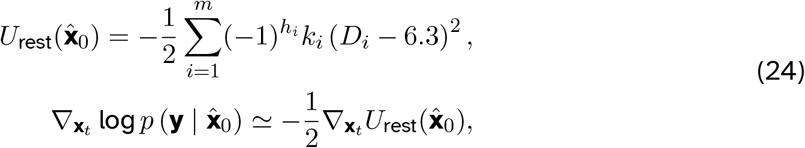

where *h*_*i*_ = 0 indicates that the *i*-th residue forms part of a helix in the sample, and *h*_*i*_ = 1 indicates that it does not. This approach ensures that the specified distance constraint is maintained, promoting helix formation in the desired region of the protein.

#### Cryo-EM image

To generate the image corresponding to a configuration, we represent the configuration with its *Cα* atoms. We first place the center of mass of the molecule at the origin, and apply a rotation matrix *R*. The electron density at location **r** = (*r*_*x*_, *r*_*y*_, *r*_*z*_) is obtained by summing over 3D Gaussian of width *σ* = 1.5Å centered at each *C*_*α*_ atoms:

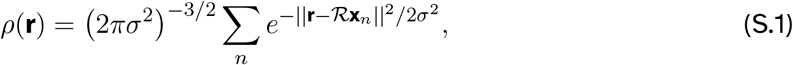

where **x**_*n*_ is the position of n-th *Cα*. The 2D intensity map *I* is obtained by projecting along the z-axis:

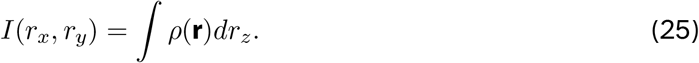

In the experiment, the 2D density map is discretized with 2D grid of size 64 * 64. Gaussian noise is added with various levels of signal-to-noise ratio.

### 5 Evaluation Metrics

#### Root Mean Squared Distance (RMSD)

Root Mean Square Deviation (RMSD) is a commonly used measure in structural biology to quantify the difference between two protein structures. It’s particularly useful for comparing the similarity of protein three-dimensional structures. The RMSD is calculated by taking the square root of the average of the square of the distances between the atoms of two superimposed proteins:

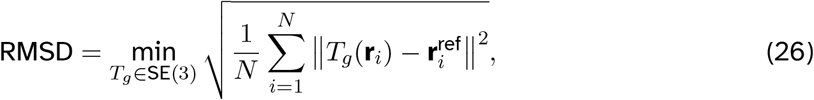

where *N* is the number of atoms in the protein, and **r**_*i*_ and 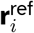 are positions of the *i*−th equivalent atoms of two structures being compared. A lower RMSD of a generated configuration indicates more similarity to the original all-atom configuration.

#### RMSF

The root-mean-square fluctuation (RMSF) is the time-averaged measure of the root-mean-square deviation (RMSD). The RMSF of a residue *ρ*_*i*_, represented by its *Cα* position **r**_*i*_, is calculated using the following equation:

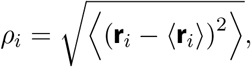

where ⟨**r**_*i*_⟩ is the ensemble average position of the *i*-th *Cα*. As with RMSD calculations, all structures are superimposed onto a reference frame configuration. Regions of the structure with high RMSF values frequently deviate from the average, indicating areas of high mobility.

#### Secondary structure

The computation of secondary structure follows Dictionary of protein secondary structure (DSSP) protocol. ^117^ The calculation is done using the Mdtraj^118^ package, and the implementation of DSSP is based on DSSP-2.2.0.

#### JS divergence

The Jensen-Shannon (JS) divergence is a method of measuring the similarity between two probability distributions. It is a symmetrized and smoothed version of the Kullback-Leibler (KL) divergence, and is always bounded between 0 and 1. For two probability distributions *P* and *Q*, the JS divergence is defined as follows:

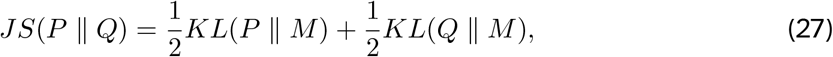

where 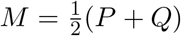 is the average of the two distributions, and *KL*(*P* ∥ *M*) is the Kullback-Leibler divergence between *P* and *M*. The KL divergence is defined as:

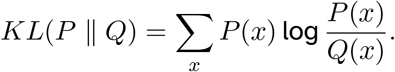

In this paper, we use JS divergence to quantify the difference in the end-to-end distance, radius of gyration, and the solvent accessible surface area.

### 6 Limitation

#### Manifold constraint sampling

One key limitation of manifold constraint sampling is the use of harmonic restraint potentials, which fail to consider the covariance between dimensions, leading to inaccuracies, especially when guiding the unified protein ensemble sampler with high-dimensional experimental data. To address this, a potential solution is the use of Mahalanobis-distance-based restraints, which inherently account for correlations between variables. This approach offers a more sophisticated method of applying restraints by measuring the distance between a point and the underlying distribution, thereby maintaining multidimensional depen-dencies and improving the fidelity of the sampling process. Another challenge arises from the reliance on automatic differentiation (autograd) during sampling. This method requires the computational graph to be retained throughout the sampling steps, resulting in higher computational costs and increased GPU usage, particularly when dealing with high-dimensional data such as 2D cryo-EM images. This bottleneck becomes more pronounced as the dimensionality of the guiding parameters increases, necessitating smaller reverse diffusion time steps to maintain accuracy, which in turn prolongs the inference time. Finally, the current implementation of ExEnDiff relies on generating or obtaining samples from experimental measurements before initiating the ensemble sampling. This approach poses a limitation if the generated samples fail to adequately represent the entire free energy surface of the measurement space. A density-guided rather than sample-guided ExEnDiff can potentially achieve much higher performance. Overcoming these limitations is crucial for advancing the manifold constraint technique and enhancing the performance of ExEnDiff, especially in addressing complex targets like cryo-EM data. By improving both the restraint model and computational efficiency, ExEnDiff can better handle the challenges of high-dimensional experimental guidance.

#### Cryo-EM

Currently, ExEnDiff uses a simplified forward configuration-to-image model to guide the sampling process. In a more complete and sophisticated model, the image *I* is convolved with the real-space point-spread function:

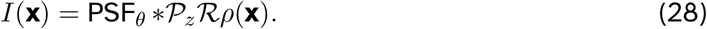

A typical Point Spread Function (PSF), as used in various previous works, ^67,119^ is given by:

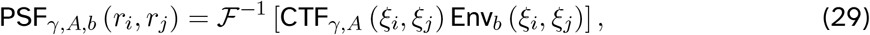

where ℱ^−1^ represents the inverse of the two-dimensional Fourier transform. The contrast transfer function (CTF) is defined as:

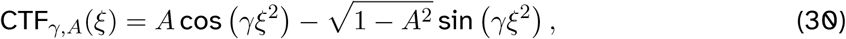

and the envelope function is:

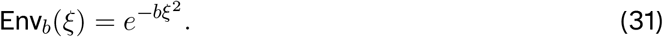

Here, *A* refers to the amplitude-contrast ratio, and *b* is the B-factor. The parameter *γ* is given by *γ* = −*π*Δ*zλ*_*e*_, where Δ*z* is the defocus and *λ*_*e*_ represents the electron wavelength. The variables *ξ*_*i*_ and *ξ*_*j*_ denote the spatial frequencies in the horizontal and vertical directions of the image, respectively.

Additionally, in the current model, the rotation matrix is pre chosen and fixed, while in a true model, the rotation matrix is drawn from the uniform distribution on SO(3). Below we formulate the process of recovering both the configuration and the unknown rotation matrix using the manifold constraint sampling technique. Since the rotation matrix and the configuration is independent, we have *p*_*t*_(**x**_*t*_, ℛ) = *p*_*t*_(**x**_*t*_)*p*(ℛ). Thus, the intermediate score function with an unknown rotation matrix can be written as:

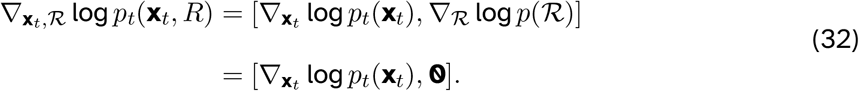

With a given 2D density Image *I*, we can rewrite the conditional intermediate score function:

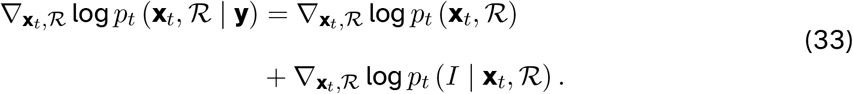

Using the manifold constraint sampling technique, the second term can be approximated as:

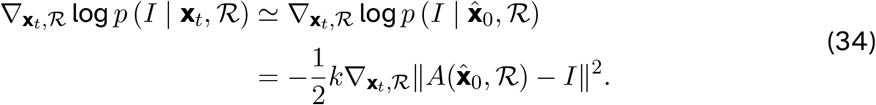

While the sampling trajectories of the configurations adhere to the manifold constraint sampling technique, the sampling of the rotation matrix follows standard overdamped Langevin dynamics. In practice, the time required for the rotation matrix to converge to the ground truth can be quite time-consuming, potentially becoming a bottleneck in the overall sampling process. Finally, adopting a Cryo-EM density image of larger sizes can be computationally difficult and resource-intensive for the current ExEnDiff model.

#### Real experimental data

In this study, we utilize synthetic experimental data that provides both the distribution of the radius of gyration (Rg) and the helix percentage for each residue in the protein. This comprehensive information allows us to accurately guide the sampling process and ensure the generated protein conformations are physically realistic. However, in real experimental scenarios, certain types of information may be incomplete or missing. For example, the SAXS database typically provides only the mean radius of gyration, without reporting its standard deviation. Additionally, detailed residue-level information such as helix percentages may not always be available, especially for flexible or disordered regions. This limitation underscores the challenge of integrating real-world experimental data, which can vary in completeness, and highlights the importance of developing methods that can operate effectively even when some measurements are missing or incomplete.

We present a potential method for estimating the standard deviation of the radius of gyration. We utilize AlphaFold3^2^ to generate the predicted template modeling (pTM) score for a set of proteins with molecular dynamics simulations results from DE Shaw Research. The pTM score reflects the confidence in AlphaFold3’s predictions, with higher scores indicating a more likely folded structure. For the proteins in the dataset, we analyzed the standard deviation of the radius of gyration and explored its relationship with the pTM score and sequence length, as shown in Fig. 3. The results show an almost linear relationship between the standard deviation of the radius of gyration and the sequence length divided by the pTM score. This observation makes intuitive sense: a higher pTM score indicates greater confidence in a properly folded protein, leading to a smaller standard deviation, while longer sequences tend to exhibit greater variance in the radius of gyration. In the future project, we aim to investigate further in how to collect low dimensional feature information from a various of experiment measurement database, computational and machine learning tools efficiently.

### 7 Additional Experimental Results

#### 7.1 Full experimental results

Here we present the experiment results of all proteins in the benchmark dataset, as demonstrated in Fig. 1 and Fig. 5. The experiment results of all benchmark proteins demonstrate the superior performance of ExEnDiff. Additionally, the comprehensive experiment results demonstrate that ExEnDiff will achieve its highest performance when guided by global features (radius of gyration) and local features (secondary structure). Furthermore, we present the mean and standard deviation of RMSF difference per residue in Fig. 7. The RMSF difference of each residue is computed as:

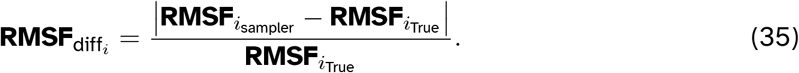

In all experiments except for RS peptide, ExEnDiff guided by Rg and the secondary structure demonstrates minimum difference in RMSF, showcasing its capacity in capturing local mobility. Finally, we present the experiment results of ExEnDiff guided by Cryo-EM 2D density images on two other proteins, as shown in Fig. 6.

#### 7.2 Choice of the correction hyperparameter

An important hyperparameter in the manifold constraint sampling is the correction term weight *k*_*i*_. We should expect that a too-low weight will lead to inconsistency with the conditions and an overly-high weight will make the sampling path noisy. Following the original manifold constraint sampling technique, ^106^ we set 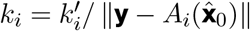. In practice, different operators can require different values of 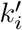. In this paper, we report the empirical values that yield the best performance in Table. 1.

#### 7.3 MESA training formulation

Autoencoder is a type of neural network used for dimensionality reduction by learning an efficient representation of data. It consists of two main parts: an encoder, which compresses the input data into a lower-dimensional latent space, and a decoder, which reconstructs the original data from this compressed representation. The network is trained to minimize the difference between the input and the reconstructed output, effectively learning key features of the data in a compact form. Autoencoder and its variants have been used extensively for learning ML-based CVs for enhanced sampling. In this work we follow MESA (Molecular enhanced sampling with autoencoders) and use the simplest form of the autoencoder. The framework is shown in Fig. 8.

The input feature is the trignometric form of dihedral angles formed by every consecutive four *Cα*. The encoder part of the model consists of a multi-layer perceptron (MLP) with the Tanh activation function, which transforms the input trignometric form of the torsion angles into a latent space of 2 dimensions: **z** = **E**_*θ*_(sin *τ*, cos *τ*). This latent space is intended to capture the essential collective variables required for describing the protein conformational landscape. The decoder, which is also an MLP using the Tanh activation function, reconstructs the input torsion angles from the latent space: *τ* ^*′*^ = **D**_*θ*_(**z**) The reconstruction loss is measured as:

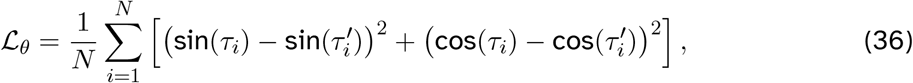

where *τ*_*i*_ and 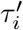 are the original and reconstructed torsion angles. The optimization is performed using the Adam optimizer with a learning rate of 0.0005, a batch size of 128, and the model is trained for 500 epochs.

## References

[1] Jumper, J.; Evans, R.; Pritzel, A.; Green, T.; Figurnov, M.; Ronneberger, O.; Tunyasuvunakool, K.; Bates, R.; Žídek, A.; Potapenko, A.; others nature 2021, 596, 583–589.

[2] Abramson, J.; Adler, J.; Dunger, J.; Evans, R.; Green, T.; Pritzel, A.; Ronneberger, O.; Willmore, L.; Ballard, A. J.; Bambrick, J.; others Nature 2024, 1–3.

[3] Baek, M.; DiMaio, F.; Anishchenko, I.; Dauparas, J.; Ovchinnikov, S.; Lee, G. R.; Wang, J.; Cong, Q.; Kinch, L. N.; Schaeffer, R. D.; others Science 2021, 373, 871–876.

[4] Du, Z.; Su, H.; Wang, W.; Ye, L.; Wei, H.; Peng, Z.; Anishchenko, I.; Baker, D.; Yang, J. Nature protocols 2021, 16, 5634–5651.

[5] Lin, Z.; Akin, H.; Rao, R.; Hie, B.; Zhu, Z.; Lu, W.; dos Santos Costa, A.; Fazel-Zarandi, M.; Sercu, T.; Candido, S.; others BioRxiv 2022, 2022, 500902.

[6] Stein, R. A.; Mchaourab, H. S. PLOS Computational Biology 2022, 18, e1010483.

[7] Wayment-Steele, H. K.; Ojoawo, A.; Otten, R.; Apitz, J. M.; Pitsawong, W.; Hömberger, M.; Ovchinnikov, S.; Colwell, L.; Kern, D. Nature 2024, 625, 832–839.

[8] Song, Y.; Sohl-Dickstein, J.; Kingma, D. P.; Kumar, A.; Ermon, S.; Poole, B. arXiv preprint 2011.13456 2020,

[9] Ho, J.; Jain, A.; Abbeel, P. Advances in neural information processing systems 2020, 33, 6840–6851.

[10] Lu, J.; Zhong, B.; Zhang, Z.; Tang, J. arXiv preprint 2306.03117 2023,

[11] Jing, B.; Erives, E.; Pao-Huang, P.; Corso, G.; Berger, B.; Jaakkola, T. arXiv preprint 2304.02198 2023,

[12] Jing, B.; Berger, B.; Jaakkola, T. arXiv preprint 2402.04845 2024,

[13] Zheng, S.; He, J.; Liu, C.; Shi, Y.; Lu, Z.; Feng, W.; Ju, F.; Wang, J.; Zhu, J.; Min, Y.; others arXiv preprint 2306.05445 2023,

[14] Ciccotti, G.; Ferrario, M.; Schuette, C.; others Entropy 2014, 16, 1.

[15] Kmiecik, S.; Gront, D.; Kolinski, M.; Wieteska, L.; Dawid, A. E.; Kolinski, A. Chemical reviews 2016, 116, 7898–7936.

[16] Marrink, S. J.; Risselada, H. J.; Yefimov, S.; Tieleman, D. P.; De Vries, A. H. The journal of physical chemistry B 2007, 111, 7812–7824.

[17] Rudd, R. E.; Broughton, J. Q. Physical review B 1998, 58, R5893.

[18] Shaw, D. E.; Maragakis, P.; Lindorff-Larsen, K.; Piana, S.; Dror, R. O.; Eastwood, M. P.; Bank, J. A.; Jumper, J. M.; Salmon, J. K.; Shan, Y.; others Science 2010, 330, 341–346.

[19] Wang, Y.; Wang, L.; Shen, Y.; Wang, Y.; Yuan, H.; Wu, Y.; Gu, Q. arXiv preprint 2403.14088 2024,

[20] Sterckx, Y. G.; Volkov, A. N.; Vranken, W. F.; Kragelj, J.; Jensen, M. R.; Buts, L.; Garcia-Pino, A.; Jové, T.; Van Melderen, L.; Blackledge, M.; others Structure 2014, 22, 854–865.

[21] Marsh, J. A.; Forman-Kay, J. D. Journal of molecular biology 2009, 391, 359–374.

[22] Ozenne, V.; Schneider, R.; Yao, M.; Huang, J.-r.; Salmon, L.; Zweckstetter, M.; Jensen, M. R.; Blackledge, M. Journal of the American Chemical Society 2012, 134, 15138–15148.

[23] Struthers, M.; Ottesen, J. J.; Imperiali, B. Folding and Design 1998, 3, 95–103.

[24] Bretscher, A.; Weber, K. Proceedings of the National Academy of Sciences 1979, 76, 2321–2325.

[25] Iesmantavicius, V.; Jensen, M. R.; Ozenne, V.; Blackledge, M.; Poulsen, F. M.; Kjaergaard, M. Journal of the American Chemical Society 2013, 135, 10155–10163.

[26] Zhang, O.; Forman-Kay, J. D. Biochemistry 1995, 34, 6784–6794.

[27] Jensen, M. R.; Communie, G.; Ribeiro Jr, E. A.; Martinez, N.; Desfosses, A.; Salmon, L.; Mollica, L.; Gabel, F.; Jamin, M.; Longhi, S.; others Proceedings of the National Academy of Sciences 2011, 108, 9839–9844.

[28] Sgourakis, N. G.; Yan, Y.; McCallum, S. A.; Wang, C.; Garcia, A. E. Journal of molecular biology 2007, 368, 1448–1457.

[29] Martin, E. W.; Holehouse, A. S.; Grace, C. R.; Hughes, A.; Pappu, R. V.; Mittag, T. Journal of the American Chemical Society 2016, 138, 15323–15335.

[30] Lindsay, R. J.; Wigge, P. A.; Hanson, S. M. BioRxiv 2023, 2023–03.

[31] Larson, S. M.; Snow, C. D.; Shirts, M.; Pande, V. S. arXiv preprint 0901.0866 2009,

[32] Lindsay, R. J.; Sahoo, A.; Liu, Y.; Hanson, S. Folding@Home simulations of RS-Peptide using Amber03ws force field and Tip4p/2005 water model. 2024; https://doi.org/10.5281/zenodo.14013285.

[33] Sahoo, H. Journal of Photochemistry and Photobiology C: Photochemistry Reviews 2011, 12, 20–30.

[34] Beljonne, D.; Curutchet, C.; Scholes, G. D.; Silbey, R. J. The journal of physical chemistry B 2009, 113, 6583–6599.

[35] de Souza, E. S.; Hirata, I. Y.; Juliano, L.; Ito, A. S. Biochimica et Biophysica Acta (BBA)-General Subjects 2000, 1474, 251–261.

[36] Möglich, A.; Joder, K.; Kiefhaber, T. Proceedings of the National Academy of Sciences 2006, 103, 12394–12399.

[37] Roy, R.; Hohng, S.; Ha, T. Nature methods 2008, 5, 507–516.

[38] Schuler, B.; Eaton, W. A. Current opinion in structural biology 2008, 18, 16–26.

[39] Hellenkamp, B.; Schmid, S.; Doroshenko, O.; Opanasyuk, O.; Kühnemuth, R.; Rezaei Adariani, S.; Ambrose, B.; Aznauryan, M.; Barth, A.; Birkedal, V.; others Nature methods 2018, 15, 669–676.

[40] Li, T.; Senesi, A. J.; Lee, B. Chemical reviews 2016, 116, 11128–11180.

[41] Chu, B.; Hsiao, B. S. Chemical reviews 2001, 101, 1727–1762.

[42] Doniach, S. Chemical Reviews 2001, 101, 1763–1778.

[43] Bernadó, P.; Mylonas, E.; Petoukhov, M. V.; Blackledge, M.; Svergun, D. I. Journal of the American Chemical Society 2007, 129, 5656–5664.

[44] Feigin, L.; Svergun, D. I.; others Structure analysis by small-angle X-ray and neutron scattering; Springer, 1987; Vol. 1.

[45] Windsor, C. G. Journal of Applied Crystallography 1988, 21, 582–588.

[46] King, S. M. Modern Techniques for Polymer Characterisation 1999, 12.

[47] Hollamby, M. J. Physical chemistry chemical physics 2013, 15, 10566–10579.

[48] Mahieu, E.; Gabel, F. Acta Crystallographica Section D: Structural Biology 2018, 74, 715–726.

[49] Woody, R. W. Methods in enzymology 1995, 246, 34–71.

[50] Berova, N.; Nakanishi, K.; Woody, R. W. Circular dichroism: principles and applications; John Wiley & Sons, 2000.

[51] Ranjbar, B.; Gill, P. Chemical biology & drug design 2009, 74, 101–120.

[52] Greenfield, N. J. Nature protocols 2006, 1, 2876–2890.

[53] Whitmore, L.; Wallace, B. A. Biopolymers: Original Research on Biomolecules 2008, 89, 392–400.

[54] Carr, H. Y.; Purcell, E. M. Physical review 1954, 94, 630.

[55] Lambert, J. B.; Mazzola, E. P.; Ridge, C. D. Nuclear magnetic resonance spectroscopy: an introduction to principles, applications, and experimental methods; John Wiley & Sons, 2019.

[56] Wüthrich, K.; Billeter, M.; Braun, W. Journal of molecular biology 1984, 180, 715–740.

[57] Camilloni, C.; De Simone, A.; Vranken, W. F.; Vendruscolo, M. Biochemistry 2012, 51, 2224–2231.

[58] Satoh, D.; Shimizu, K.; Nakamura, S.; Terada, T. FEBS letters 2006, 580, 3422–3426.

[59] Maruyama, Y.; Koroku, S.; Imai, M.; Takeuchi, K.; Mitsutake, A. RSC advances 2020, 10, 22797–22808.

[60] Airas, J.; Ding, X.; Zhang, B. ACS Central Science 2023, 9, 2286–2297.

[61] Bonomi, M.; Vendruscolo, M. Current Opinion in Structural Biology 2019, 56, 37–45.

[62] Bai, X.-C.; McMullan, G.; Scheres, S. H. Trends in biochemical sciences 2015, 40, 49–57.

[63] Cheng, Y. Cell 2015, 161, 450–457.

[64] heng, Y. Science 2018, 361, 876–880.

[65] Wrapp, D.; Wang, N.; Corbett, K. S.; Goldsmith, J. A.; Hsieh, C.-L.; Abiona, O.; Graham, B. S.; McLellan, J. S. Science 2020, 367, 1260–1263.

[66] Terwilliger, T. C.; Ludtke, S. J.; Read, R. J.; Adams, P. D.; Afonine, P. V. Nature methods 2020, 17, 923–927.

[67] Tang, W. S.; Silva-Sánchez, D.; Giraldo-Barreto, J.; Carpenter, B.; Hanson, S. M.; Barnett, A. H.; Thiede, E. H.; Cossio, P. The Journal of Physical Chemistry B 2023, 127, 5410–5421.

[68] Trabuco, L. G.; Villa, E.; Schreiner, E.; Harrison, C. B.; Schulten, K. Methods 2009, 49, 174–180.

[69] Igaev, M.; Kutzner, C.; Bock, L. V.; Vaiana, A. C.; Grubmüller, H. Elife 2019, 8, e43542.

[70] Mori, T.; Terashi, G.; Matsuoka, D.; Kihara, D.; Sugita, Y. Journal of chemical information and modeling 2021, 61, 3516–3528.

[71] Blau, C.; Yvonnesdotter, L.; Lindahl, E. PLOS Computational Biology 2023, 19, e1011255.

[72] Vuillemot, R.; Miyashita, O.; Tama, F.; Rouiller, I.; Jonic, S. Journal of Molecular Biology 2022, 434, 167483.

[73] Levy, A.; Chan, E. R.; Fridovich-Keil, S.; Poitevin, F.; Zhong, E. D.; Wetzstein, G. arXiv preprint 2406.04239 2024,

[74] Das, P.; Moll, M.; Stamati, H.; Kavraki, L. E.; Clementi, C. Proc. Natl. Acad. Sci. U.S.A. 2006, 103, 9885–9890.

[75] Ferguson, A. L.; Panagiotopoulos, A. Z.; Debenedetti, P. G.; Kevrekidis, I. G. J. Chem. Phys. 2011, 134, 135103.

[76] Rohrdanz, M. A.; Zheng, W.; Maggioni, M.; Clementi, C. J. Chem. Phys. 2011, 134, 124116.

[77] Noé, F.; Clementi, C. J. Chem. Theory Comput 2015, 11, 5002–5011.

[78] Zhang, J.; Chen, M. Physical review letters 2018, 121, 010601.

[79] Fu, X.; Xie, T.; Rebello, N. J.; Olsen, B. D.; Jaakkola, T. arXiv preprint 2204.10348 2022,

[80] Ceriotti, M.; Tribello, G. A.; Parrinello, M. Proc. Natl. Acad. Sci. U.S.A. 2011, 108, 13023–13028.

[81] Tiwary, P.; Berne, B. Proc. Natl. Acad. Sci. U.S.A. 2016, 113, 2839–2844.

[82] Mendels, D.; Piccini, G.; Parrinello, M. J. Phys. Chem. Lett. 2018, 9, 2776–2781.

[83] Ribeiro, J. M. L.; Bravo, P.; Wang, Y.; Tiwary, P. J. Chem. Phys. 2018, 149, 072301.

[84] Wang, Y.; Ribeiro, J. M. L.; Tiwary, P. Nat. Commun. 2019, 10, 3573.

[85] Nuske, F.; Keller, B. G.; Pérez-Hernández, G.; Mey, A. S.; Noé, F. J. Chem. Theory Comput 2014, 10, 1739–1752.

[86] Vani, B. P.; Aranganathan, A.; Wang, D.; Tiwary, P. Journal of chemical theory and computation 2023, 19, 4351–4354.

[87] Chen, W.; Tan, A. R.; Ferguson, A. L. The Journal of chemical physics 2018, 149.

[88] Robustelli, P.; Kohlhoff, K.; Cavalli, A.; Vendruscolo, M. Structure 2010, 18, 923–933.

[89] Leelananda, S. P.; Lindert, S. J. Chem. Inf. Model. 2020, 60, 2522–2532.

[90] Chen, P.-c.; Shevchuk, R.; Strnad, F. M.; Lorenz, C.; Karge, L.; Gilles, R.; Stadler, A. M.; Hennig, J.; Hub, J. S. J. Chem. Theory Comput. 2019, 15, 4687–4698.

[91] Roux, B.; Islam, S. M. J. Phys. Chem. B 2013, 117, 4733–4739.

[92] Dimura, M.; Peulen, T.-O.; Sanabria, H.; Rodnin, D.; Hemmen, K.; Hanke, C. A.; Seidel, C. A. M.; Gohlke, H. Nat Commun 2020, 11, 5394.

[93] Sahoo, A.; Lee, P.-Y.; Matysiak, S. J. Chem. Theory Comput. 2022, 18, 5046–5055.

[94] Arts, M.; Garcia Satorras, V.; Huang, C.-W.; Zugner, D.; Federici, M.; Clementi, C.; Noé, F.; Pinsler, R.; van den Berg, R. Journal of Chemical Theory and Computation 2023, 19, 6151–6159.

[95] Liu, Y.; Ghosh, T. K.; Lin, G.; Chen, M. The Journal of Physical Chemistry Letters 2024, 15, 3938–3945.

[96] Giraldo-Barreto, J.; Ortiz, S.; Thiede, E. H.; Palacio-Rodriguez, K.; Carpenter, B.; Barnett, A. H.; Cossio, P. Scientific reports 2021, 11, 13657.

[97] Bradshaw, R. T.; Marinelli, F.; Faraldo-Gómez, J. D.; Forrest, L. R. Biophysical Journal 2020, 118, 1649–1664.

[98] Hummer, G.; Köfinger, J. The Journal of Chemical Physics 2015, 143, 243150.

[99] Różycki, B.; Kim, Y. C.; Hummer, G. Structure 2011, 19, 109–116.

[100] Reichel, K.; Stelzl, L. S.; Köfinger, J.; Hummer, G. J. Phys. Chem. Lett. 2018, 9, 5748–5752.

[101] Köfinger, J.; Stelzl, L. S.; Reuter, K.; Allande, C.; Reichel, K.; Hummer, G. J. Chem. Theory Comput. 2019, 15, 3390–3401.

[102] Feigin, L. A.; Svergun, D. I. In Structure Analysis by Small-Angle X-Ray and Neutron Scattering; Taylor, G. W., Ed.; Springer US: Boston, MA, 1987.

[103] Jacques, D. A.; Trewhella, J. Protein Science 2010, 19, 642–657.

[104] Putnam, C. D.; Hammel, M.; Hura, G. L.; Tainer, J. A. Quart. Rev. Biophys. 2007, 40, 191–285.

[105] Anderson, B. D. Stochastic Processes and their Applications 1982, 12, 313–326.

[106] Chung, H.; Kim, J.; Mccann, M. T.; Klasky, M. L.; Ye, J. C. arXiv preprint 2209.14687 2022,

[107] Robustelli, P.; Piana, S.; Shaw, D. E. Proceedings of the National Academy of Sciences 2018, 115, E4758–E4766.

[108] Lindorff-Larsen, K.; Piana, S.; Dror, R. O.; Shaw, D. E. Science 2011, 334, 517–520.

[109] Piana, S.; Lindorff-Larsen, K.; Shaw, D. E. Proceedings of the National Academy of Sciences 2013, 110, 5915–5920.

[110] Mark, P.; Nilsson, L. The Journal of Physical Chemistry A 2001, 105, 9954–9960.

[111] Brooks, B. R.; Brooks III, C. L.; Mackerell Jr, A. D.; Nilsson, L.; Petrella, R. J.; Roux, B.; Won, Y.; Archontis, G.; Bartels, C.; Boresch, S.; others Journal of computational chemistry 2009, 30, 1545–1614.

[112] Wang, L.; Friesner, R. A.; Berne, B. The Journal of Physical Chemistry B 2011, 115, 9431–9438.

[113] Lindsay, R. J.; Viegas, R. G.; Leite, V. B.; Wigge, P. A.; Hanson, S. M. bioRxiv 2024, 2024–08.

[114] Wang, J.; Wolf, R. M.; Caldwell, J. W.; Kollman, P. A.; Case, D. A. Journal of computational chemistry 2004, 25, 1157–1174.

[115] Abascal, J. L.; Vega, C. The Journal of chemical physics 2005, 123.

[116] Hess, B.; Bekker, H.; Berendsen, H. J.; Fraaije, J. G. Journal of computational chemistry 1997, 18, 1463–1472.

[117] Kabsch, W.; Sander, C. Biopolymers: Original Research on Biomolecules 1983, 22, 2577–2637.

[118] McGibbon, R. T.; Beauchamp, K. A.; Harrigan, M. P.; Klein, C.; Swails, J. M.; Hernández, C. X.; Schwantes, C. R.; Wang, L.-P.; Lane, T. J.; Pande, V. S. Biophysical journal 2015, 109, 1528–1532.

[119] Cossio, P.; Hummer, G. Journal of structural biology 2013, 184, 427–437.

